# Integrative transcriptome imputation reveals tissue-specific and shared biological mechanisms mediating susceptibility to complex traits

**DOI:** 10.1101/532929

**Authors:** Wen Zhang, Georgios Voloudakis, Veera M. Rajagopal, Ben Reahead, Joel T. Dudley, Eric E. Schadt, Johan L.M. Björkegren, Yungil Kim, John F. Fullard, Gabriel E. Hoffman, Panos Roussos

## Abstract

Transcriptome-wide association studies integrate gene expression data with common risk variation to identify gene-trait associations. By incorporating epigenome data to estimate the functional importance of genetic variation on gene expression, we improve the accuracy of transcriptome prediction and the power to detect significant expression-trait associations. Joint analysis of 14 large-scale transcriptome datasets and 58 traits identify 13,724 significant expression-trait associations that converge to biological processes and relevant phenotypes in human and mouse phenotype databases. We perform drug repurposing analysis and identify known and novel compounds that mimic or reverse trait-specific changes. We identify genes that exhibit agonistic pleiotropy for genetically correlated traits that converge on shared biological pathways and elucidate distinct processes in disease etiopathogenesis. Overall, this comprehensive analysis provides insight into the specificity and convergence of gene expression on susceptibility to complex traits.

---

Despite the recent success of genome-wide association studies (GWASs) in cataloguing risk genetic variation, our understanding of the mechanisms through which they act remain largely unknown^1^. Risk variants are highly enriched in *cis* regulatory elements (CREs), including promoters and enhancers^2,3^ and affect the regulation of gene expression^2–10^. Multiple computational methods have been developed to link risk variants with differential gene expression^11–15^. PrediXcan^16^ performs transcriptome-wide association study (TWAS) by gene expression imputation, and so far it outperforms similar methods^17^. Briefly, PrediXcan uses elastic net (ENet) regression models, trained in a reference transcriptome, to impute gene expression. The models use a set of *cis*-SNPs (SNPs in proximity to the transcription start site) as linear predictors of gene expression. The imputed expressions are then correlated with the phenotype of interest to identify gene-trait associations (GTAs).

Here we present EpiXcan, a novel method that increases prediction accuracy in transcriptome imputation by integrating epigenetic data to model the prior probability that a SNP affects transcription. EpiXcan specifically leverages annotations derived from the Roadmap Epigenomics Mapping Consortium (REMC) that integrates multiple epigenetic assays, including DNA methylation, histone modification and chromatin accessibility^18^. The rationale of our approach is that SNPs within CREs are more likely to be functionally relevant^19^. We then utilize 14 large-scale transcriptome datasets of genotyped individuals to train prediction models and integrate with 58 complex traits and diseases to define significant GTAs. GTAs exhibit significant enrichment for relevant biological pathways and known genes linked to trait-related phenotypes in humans and mice. Imputed transcriptomic changes are used to identify known compounds that can normalize genetically driven expression perturbations. Pairwise trait analysis identifies genes that exhibit agonistic pleiotropy for genetically correlated traits that converge on shared biological pathways. Finally, bi-directional regression analysis identifies putative causal relationships among traits. Overall, our analysis provides insight into the specificity and convergence of gene expression mediating the genetic risk architecture underlying susceptibility of complex traits and diseases.

## Results

### EpiXcan outperforms PrediXcan

Since TWAS is limited to genes that can be accurately predicted from genotype data, increasing prediction accuracy can increase the scope and power of analyses. Here, we integrate biologically relevant data in a single framework to improve performance of gene expression prediction. The overall schematic of EpiXcan is shown in **Supplementary Fig. 1**. Briefly, EpiXcan leverages epigenetic annotation to inform transcriptomic imputation by employing a three-step process (**Online methods** and **Supplementary Methods**): *(1)* estimate SNP priors that reflect the likelihood of a SNP having a regulatory role in gene expression based on a Bayesian hierarchical model^20^ that integrates epigenomic annotation^18^ and eQTL summary statistics for *cis*-SNPs (SNPs located ± 1 Mb from the transcription start site of the gene); *(2)* rescale the SNP priors to penalty factors by employing a novel adaptive mapping approach; and *(3)* use the genotypes and penalty factors in weighted elastic net to perform gene expression prediction.

Using simulated data, we apply EpiXcan and PrediXcan to train prediction models and estimate the adjusted cross-validation R-squared (*R^2^ _CV_*), which is the correlation between the predicted and observed expression levels during the nested cross validation. In all simulated scenarios, EpiXcan improves the average *R^2^_CV_* compared to PrediXcan models (all *p* values ≤ 7 × 10^-10^ based on one-sample sign test; **Supplementary Fig. 2**). We then train prediction models by applying EpiXcan and PrediXcan in 14 RNAseq datasets, derived from dorsolateral prefrontal cortex (DLPFC) from the CommonMind Consortium (CMC)^21^, seven tissues from Stockholm-Tartu Atherosclerosis Reverse Network Engineering Task (STARNET)^22^ and six tissues from GTEx^23^ (**Supplementary Table 1**). We compare the performance of EpiXcan with PrediXcan models, by considering the delta value (EpiXcan minus PrediXcan) of two metrics: *(1)* crossvalidation *R^2^* (*R^2^_CV_*) within each tissue and *(2)* predictive performance *R^2^*(*R^2^_PP_*) estimated based on Pearson’s correlation between predicted and observed expression in an independent dataset of a relevant tissue. Positive delta values indicate that EpiXcan has higher prediction performance compared to PrediXcan.

Across all datasets, EpiXcan improves the average *R^2^_CV_* compared to PrediXcan (all *p* values ≤ 9 × 10^-16^ based on one-sample sign test; **Fig. 1; Supplementary Fig. 3; Supplementary Table 2**). We predict 4.6% more genes (pairwise Wilcoxon test *p* value = 6.10 × 10^-5^) with *R^2^_CV_* > 0.01 using EpiXcan (average number of genes across tissues is 10,181) compared to PrediXcan (average number of genes across tissues is 9,760). To obtain the second metric, *R^2^_PP_*, we train prediction models in the training dataset, which are then used to predict expressions in the test dataset. Across all datasets, EpiXcan improves the average *R^2^_PP_* compared to PrediXcan (all *p* values < 9 × 10^-16^ based on one-sample sign test; **Fig. 1; Supplementary Fig. 4; Supplementary Table 3**). Importantly, the ratios of genes predicted more effectively by EpiXcan than PrediXcan are higher in the independent dataset evaluation (*R^2^PP*) than in the cross-validation (unpaired t-test, *p* value = 3.3 × 10^-17^) (**Fig. 1**), suggesting that the adaptive rescaling of the penalty factors during model training does not result in significant overfitting that could affect the external validity of the models. Overall, compared to PrediXcan, EpiXcan has improved predictive performance and identifies more genes that can be used for TWAS.

**Figure 1.**
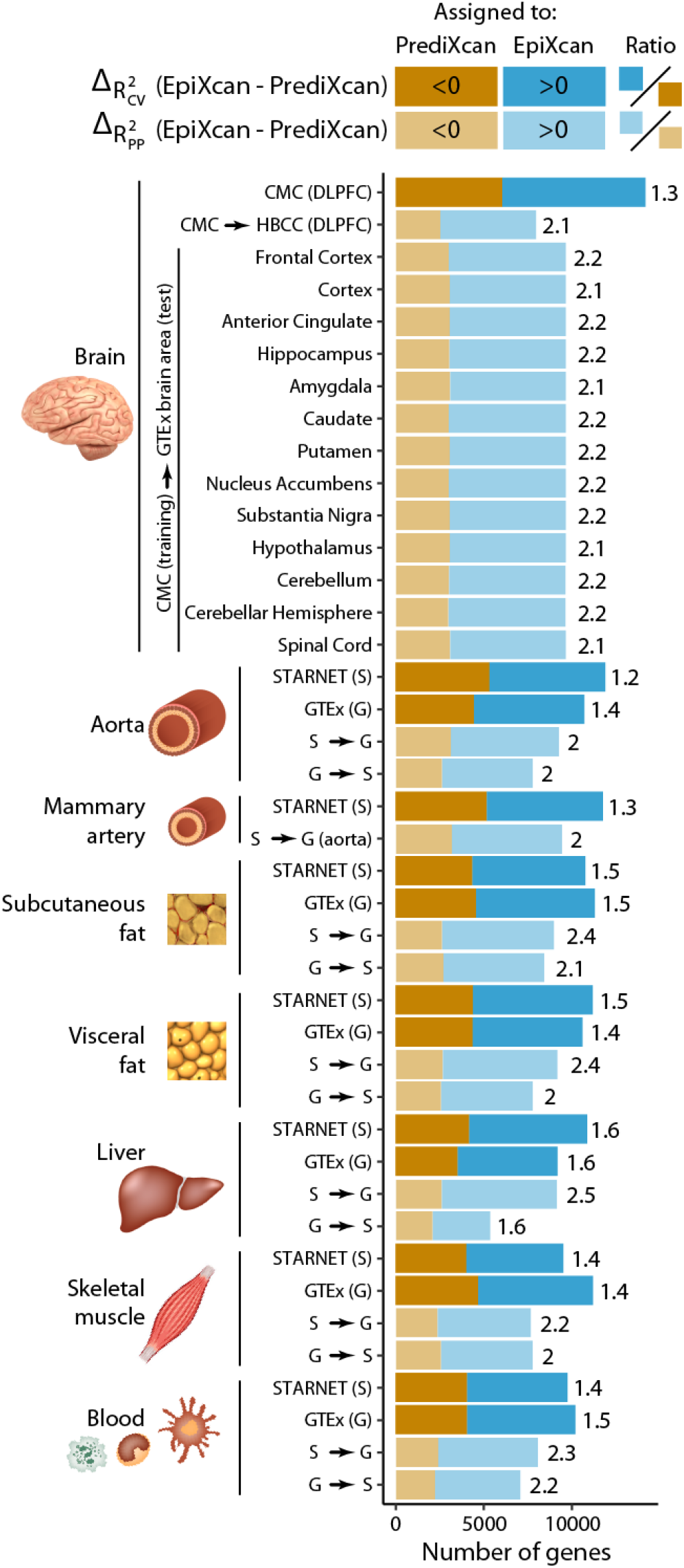
Comparison of prediction performance between EpiXcan and PrediXcan. EpiXcan and PrediXcan models are trained across multiple tissues that include: brain, aorta, mammary artery, subcutaneous fat, visceral fat, liver, skeletal muscle and blood by leveraging 14 datasets from CMC, STARNET and GTEx. The difference in training performance between EpiXcan and PrediXcan models is compared using the adjusted cross validation R^2^ (*R^2^_CV_*) metric. The 14 models are further assessed by estimating the predictive performance (*R^2^_PP_*) in independent datasets; the training dataset is shown before the arrow and the test dataset after the arrow (G = GTEx and S = STARNET). For a given dataset, we compare the *R^2^_CV_* and *R_PP_* by estimating the delta value of EpiXcan minus PrediXcan for each gene. Positive and negative delta values indicate genes with higher predictive performance in EpiXcan and PrediXcan, respectively. These genes are assigned as “EpiXcan” and “PrediXcan” and counts are shown as barplots. The number on the right indicates the ratio of “EpiXcan” assigned gene counts divided by “PrediXcan” counts. Across all datasets, the ratios are higher than 1 indicating that EpiXcan outperforms PrediXcan. *p* value from one-sample sign test indicates that the shift of the delta *R^2^_CV_* and *R_PP_* values is greater than zero (All *p* values < 9 × 10^-16^).

### EpiXcan informs better gene-trait associations than PrediXcan

We apply EpiXcan and PrediXcan prediction models from 14 tissues (**Supplementary Table 1**) in 58 complex traits (**Supplementary Table 4**) and examine their performance based on four criteria: the number of GTAs that are: *(1)* significant after multiple testing correction, *(2)* novel (significantly associated genes that lie outside the GWAS loci) *(3)* unique (i.e., genes identified only by one method), and *(4)* enriched for clinically relevant genes.

EpiXcan has more power to detect GTAs than PrediXcan (Kolmogorov-Smirnov *p* value is 3.3 × 10^-16^; **Fig. 2a**). We observe a 9.6% increase (n = 1,202) in the significant GTAs at 0.01 false discovery rate (FDR)^24^ using EpiXcan (n=13,724) compared to PrediXcan (n=12,522). One advantage of PrediXcan/EpiXcan methods is that they identify “novel genes” within loci that did not reach genome-wide significance (*p* < 5 × 10^-8^) in GWASs. We detect an 18.3% increase (one sample sign test *p* value = 3.6 × 10^-6^) in the novel GTAs using EpiXcan (mean of 25.4) compared to PrediXcan (mean of 21.5) (**Supplementary Fig. 5**). The largest difference is observed for height (EpiXcan = 168, PrediXcan = 134), followed by schizophrenia (EpiXcan = 119, PrediXcan = 104) (**Supplementary Fig. 6**).

**Figure 2.**
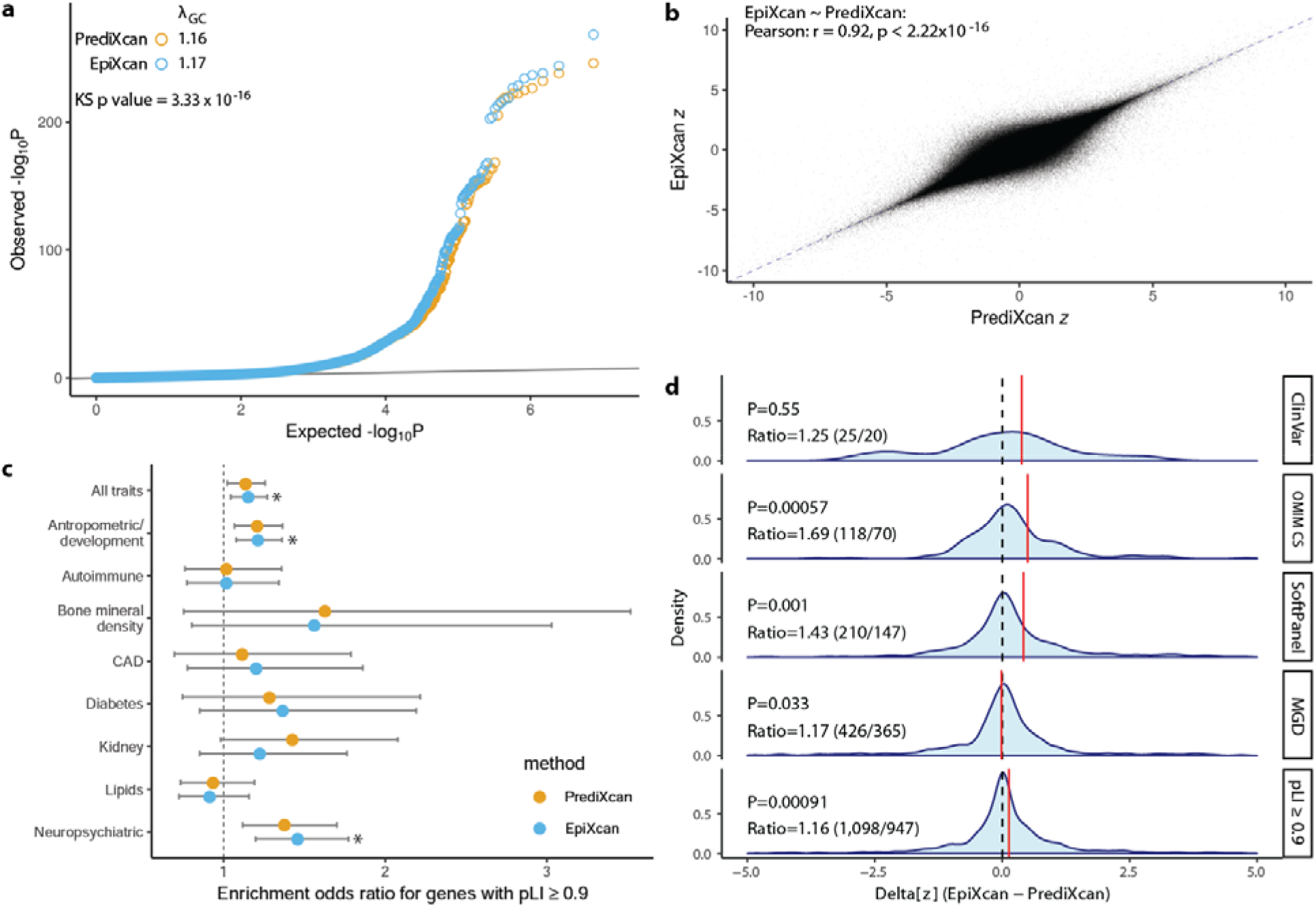
Comparison of gene-trait associations between EpiXcan and PrediXcan. **(a)** EpiXcan and PrediXcan *p* value distributions for all gene-trait associations. Quantile-quantile (QQ) plot of the *p* values for all gene-trait associations show a shift to the left. The genomic inflation factor (λ) is slightly higher for EpiXcan than PrediXcan (1.17 and 1.16). The two distributions are significantly different (Kolmogorov-Smirnov test *p* value is 3.3 × 10^-16^). **(b)** EpiXcan and PrediXcan have a high correlation of gene-trait association z-scores. Scatter plot of EpiXcan and PrediXcan Z values, Pearson r = 0.92 and Spearman *ρ* = 0.91, *p* value < 2.22 × 10^-16^ for both. Only z values between −10 and 10 are plotted. The dotted blue line corresponds to *y* = *x*. **(c)** Gene set enrichment analysis (GSEA) for extremely loss-of-function intolerant (pLI ≥ 0.9) genes. Odds ratio with 95% CI are plotted for combined gene-trait associations from all traits and trait categories for enrichment in genes with pLI ≥ 0.9 (* for *q* value <0.05). For all pLI decile bins enrichment refer to **Supplementary Table 5. (d)** EpiXcan has more power than PrediXcan to detect expression changes of trait-specific, clinically significant genes. These density plots depict the distribution of the Δ[z] (EpiXcan – PrediXcan) values for all gene-trait associations that are significant from either EpiXcan or PrediXcan. *P* value is from one sample sign test. Ratio is the number of Δ[z] measurements in favor of EpiXcan to that of PrediXcan. The red lines correspond to the mean of each distribution.

For any given tissue and trait, we find high correlation of GTA z-scores between EpiXcan and PrediXcan (Pearson’s correlation *r = 0.92*) (**Fig. 2b**) but overall, we observe unique associations for each method. We identify 79.9% (n=327) more unique genes in EpiXcan (n=788) than PrediXcan (n=461) (**Supplementary Fig. 7**), due to either a lack of a prediction model for a specific gene and/or tissue or insufficient statistical power using PrediXcan models. For example, using the waist-adjusted BMI trait and prediction models from STARNET subcutaneous adipose tissue, overall, we observe high correlation between EpiXcan and PrediXcan genes (Pearson’s *r* = 0.83) (**Supplementary Fig. 8**). Interestingly, EpiXcan identifies 7 genes (*PPP2R5A, ALAS1, HOXC8, PIEZO1, SCD, PARP3, EYA1*) that are not detected by PrediXcan even if we test across all tissue-specific models. *SCD* (stearoyl–CoA desaturase) is of particular interest as it codes for an enzyme that catalyzes a rate-limiting step in the synthesis of unsaturated fatty acids (mainly oleate and palmitoleate); knocking out the *SCD* mouse ortholog gene results in reduced body adiposity and resistance to diet-induced weight gain^25^. Accordingly, EpiXcan predicts that up-regulated *SCD* gene expression is associated with increased waist-adjusted BMI.

### EpiXcan uncovers more clinically relevant genes and molecular pathways

We perform a series of gene set enrichment analyses (GSEA) to determine how well EpiXcan can uncover clinically relevant genes and molecular pathways compared to PrediXcan. For this, we employ five categories of datasets: *(1)* ExAC gene pLI (probability of loss-of-function intolerance) dataset^26^, *(2)* ClinVar dataset - pathogenic or likely pathogenic genes in the ClinVar database^27^, *(3)* OMIM CS dataset - genes in OMIM with phenotypes in the clinical synopsis (CS) section^28^, *(4)* SoftPanel dataset - custom gene panels for our traits created with SoftPanel^29^ based on ICD-10 classification and keyword queries (underlying knowledge base is OMIM but gene panel creation is more integrative), and *(5)* MGD dataset - ortholog human genes of mouse genes associated with mouse strain-specific phenotypes^30^. GTAs from both PrediXcan and EpiXcan exhibit enrichment for genes that are associated with the traits in above datasets (**Supplementary Fig. 9**).

Transcripts identified by EpiXcan (*q* value = 0.029), but not by PrediXcan (*q* value = 0.096), are enriched for genes that are extremely loss-of-function intolerant (pLI ≥ 0.9) (**Fig. 2c**). More specifically, we find significant enrichment of pLI genes with neuropsychiatric (**q** value = 0.012, known association^31,32^) and anthropometric/development (*q* value = 0.032) related traits (**Supplementary Table 5**). Unlike pLI, for all other gene sets (ClinVar, OMIC CS, SoftPanel, MGD), we define and test for enrichment only for that specific trait. For example, for autism, we generate a gene list from the significant autism-specific GTAs from all tissues for each method and perform GSEA for genes in the ClinVar database that are reported to be associated with autism. In so doing, we find that, overall, EpiXcan has more power than PrediXcan to identify clinically relevant genes (**Fig. 2d**) including those that are more likely to belong to more than one dataset (pLI, ClinVar, OMIC CS, SoftPanel, MGD) (**Supplementary Fig. 10**).

In conclusion, TWAS across 58 traits shows that, compared to PrediXcan, EpiXcan has more power to detect significant genes, including novel and unique associations, which are indispensable for life and clinically significant. In the following section, we further explore the EpiXcan-derived GTAs, in terms of: *(1)* per-tissue contribution of significant genes, *(2)* gene-set enrichment analysis, *(3)* computational drug repurposing analysis, and *(4)* genes shared within and across different disease categories.

### Contribution of Tissues to the Identification of Associated Genes

In this study we employ 3 different training cohorts to generate 14 predictive models for 8 tissue homogenate types and use the predictive models to impute tissue-specific transcriptomes across 58 GWASs. We first determine the robustness of our method, by examining the z-score correlation for similar tissues within and across cohorts by pooling together imputed transcriptomes for each tissue from all traits. As expected, predictions are highly correlated when EpiXcan models are trained in *(1)* different cohorts (GTEx and STARNET) predicting the same tissue (Spearman’s *ρ*: 0.89-0.93) and *(2)* the same cohort predicting similar tissues (Spearman’s *ρ*: 0.89 when comparing aorta with mammary artery, and 0.92-0.95 when comparing visceral with subcutaneous adipose tissues) (**Fig. 3a**). In contrast, unrelated tissues such as blood and brain only exhibit moderate correlation (Spearman’s *ρ* 0.38 - 0.42).

**Figure 3.**
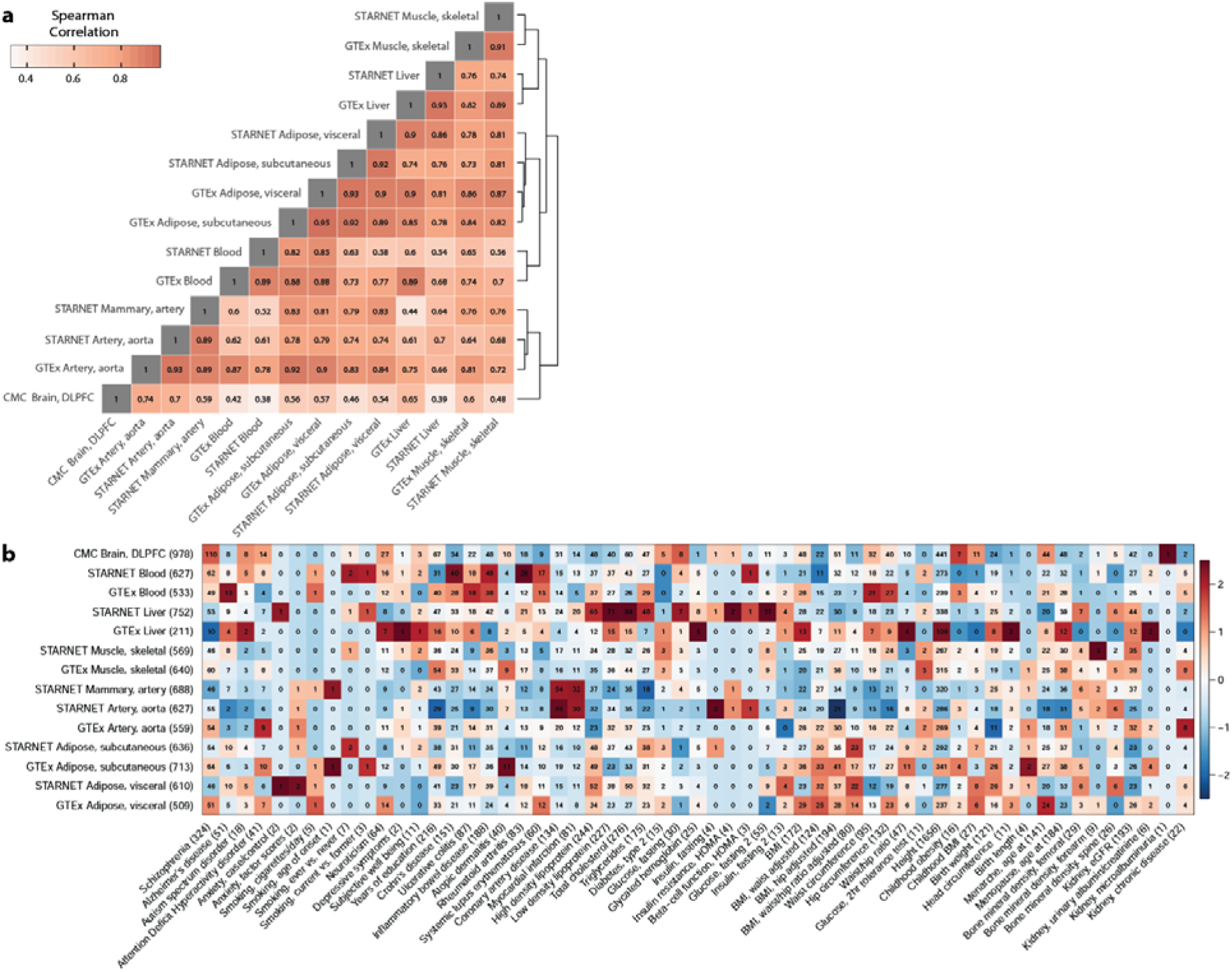
Contribution of GWAS and tissues to gene-trait associations. **(a)** Correlation of genetically regulated expression imputed for different tissues (pooled GTAs for all traits). Correlation is calculated for significant imputed expression changes with the Spearman method. Dendrogram on the right edge is shown from Ward hierarchical clustering. **(b)** Gene contribution and enrichment of each tissue prediction model for each trait. Digits in the matrix correspond to the number of genes that exhibit significant association with the trait for a given tissue prediction model. The color indicates enrichment (red) or depletion (blue) of correlations for a given trait per tissue. Numbers in parentheses alongside labels denote how many unique genes are identified from each dataset (row annotation on the left) or how many unique genes are associated with each of the traits (column annotation on the bottom).

Overall, per tissue, the numbers are largely comparable between consortia (**Supplementary Fig. 11**). However, trait-relevant tissue models contribute disproportionately more gene-trait associations than non-relevant tissues (**Fig. 3b**). For example, we find a higher number of contributions than the average of other tissues from brain tissue in schizophrenia, from arterial tissues in cardiovascular disease, and from liver in lipid traits, respectively, which is concordant with previous Summary-data-based Mendelian Randomization analysis^6^. For 48 of the traits, more than 50% of the associated genes are only found in one tissue (**Supplementary Fig. 12**) and a large proportion (32.98% ± 17.36%; mean ± SD) of these unique GTAs come from the highest contributing tissue type (**Supplementary Fig. 13**). A few examples of top tissue type contributors for unique GTAs are as follows; schizophrenia: brain tissue (30.34%, CMC), myocardial infarction & coronary artery disease: arterial tissue (33.33% & 31.88% respectively, STARNET aorta and mammary artery and GTEx aorta), systemic lupus erythematosus: blood (38.89%, STARNET and GTEx blood), most lipid traits: liver (24.06% - 26.43%, STARNET and GTEx liver). Besides tissue relevance, cohort size and tissue dissimilarity explain 52% of the variation in the number of unique GTAs contributed by different tissues (**Supplementary Fig. 14**, multiple linear regression model, *p* value = 0.007), indicating that additional GTAs will be uncovered with increased sample size of gene expression datasets in disease-relevant tissues.

### Biological relevance of gene-trait associations

GTAs for a given trait are enriched for genes implicated in diseases with more severe trait-specific phenotypes driven by larger effect mutations in those genes (ClinVar: *λ* = 1.83, *p* value = 7.07 × 10^-14^; SoftPanel: *λ* = 1.36, *p* value < 2.22 × 10^-16^; OMIM CS: *λ* = 1.29, *p* value = 6.17 × 10^-14^; **Supplementary Fig. 15**). We also observe enrichment (*λ* = 1.21, *p* value = 1.69 × 10^-13^) for genes that produce mouse phenotypes in the same phenotypic category of the human trait when the mouse ortholog gene is disrupted.

We perform gene-set enrichment analysis for the traits with more than 10 significant GTAs (43 out of 58 traits) to determine if the associated genes can be mapped to biological processes (**Supplementary Table 6**). After FDR adjustment, 74 highly enriched pathways are obtained with *p* values < 1.70 × 10^-5^ (corresponds to *q* < 0.05). Significantly associated genes are enriched for biological processes relevant to trait pathophysiology. For instance, the enriched pathways for elevated total cholesterol and triglycerides are involved in sterol and lipid homeostasis as well as lipoprotein digestion, mobilization, and transport. Similarly, for atopic dermatitis, the significantly enriched pathway modulates the rate or extent of water loss from an organism via the skin. Genes associated with mineral density of the femoral bone demonstrate a high enrichment for a pathway that positively regulates cartilage development.

### Leveraging gene-trait associations for computational drug repurposing

Computational drug repurposing offers a systematic approach for relating disease and drug-induced states towards the goal of identifying novel indications for existing therapeutics^33^. We perform a computational screen against a library of 1,309 drug-induced transcriptional profiles^34^ to identify small molecules capable of perturbing the expression of our identified trait-associated genes (**Fig. 4a**). For each trait /compound pair, we calculate a signed “connectivity score”^34^, which summarizes the transcriptional relationship between each trait and drug signature, thus identifying drugs that might be predicted to “normalize” the gene-trait signature, as well as those expected to induce a “disease-like” state (**Fig. 4b-d, Supplementary Table 7**). **Fig. 4e** provides example compounds predicted to regulate the expression of genes associated with the “Hip circumference adjusted BMI” trait. This list includes drugs under investigation for treatment of obesity, including ursolic acid, which is reported to increase skeletal muscle and brown fat while reducing diet induced obesity^35^.

**Figure 4.**
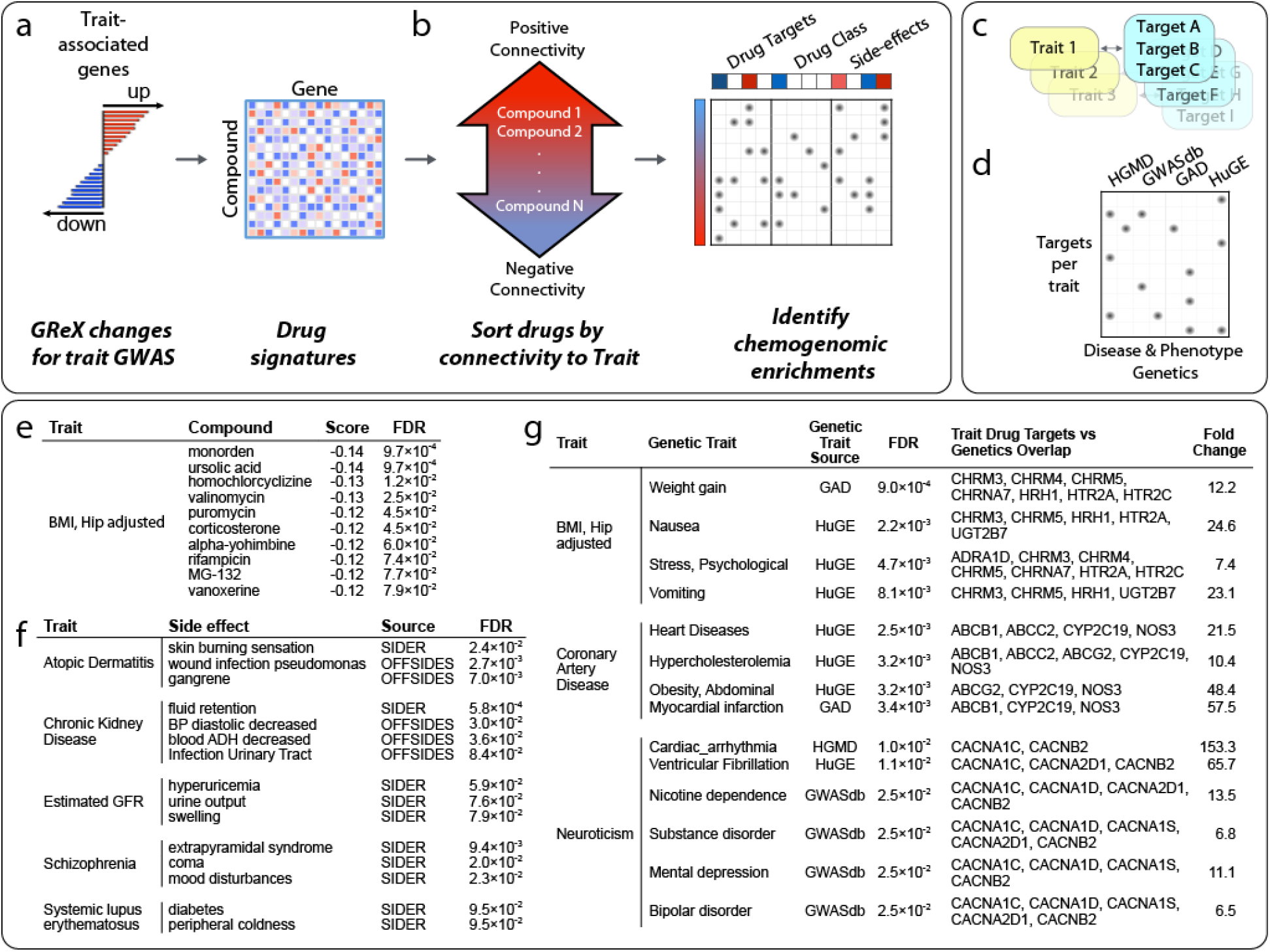
Leveraging gene-trait associations for computational drug repurposing. **(a)** Trait-associated genes are used to sort a library of drug induced gene expression signatures according to their connectivity with the trait. GReX: genetically regulated expression **(b)** A secondary enrichment analysis on this drug list identifies pharmacological features that are over-represented at the extreme ends of the sorted list, thus presenting a chemogenomic view of the trait. **(c)** Drug targets linked with each trait (FDR < 0.1) are then (d) compared with risk loci genes for a range of diseases or phenotypes (FDR < 0. 1). **(e)** Top 10 compounds predicted to normalize the expression of “Hip adjusted BMI” associated genes. **(f)** Subset of side-effect enrichments for phenotypically related traits. **(g)** Subset of traits with associated drug targets that are enriched for risk associated genes sets with phenotypically related traits.

To explore the higher-level biological context for trait/compound associations, we perform a chemogenomic enrichment analysis to determine whether drugs that regulate particular sets of trait-associated genes might share pharmacological features, such as drug targets, drug classes and side-effects (**Fig. 4b**). We find multiple significant (FDR < 0.1) chemogenomic trends, including enrichments with phenotypically related side-effects (**Fig. 4f**), supporting the potential for these compounds to perturb trait-related molecular networks.

We hypothesized that, in general, trait-associated drug targets would connect to risk-associated genes for phenotypically related diseases^36^. To evaluate this, we identify referenced^37^ and predicted^38^ drug targets that are enriched (FDR < 0.1) among compounds that modulate the signature of each trait. We identify ≥ 1 drug target enrichment, for 53 of the traits considered, and ≥ 3 drug targets for 40 traits (**Supplementary Table 7**). We then perform a further gene set analysis on the targets associated with each trait, focusing on disease risk genetic resources that might implicate phenotypes that could then be related to the traits considered within this study. We identify several significant overlaps (FDR < 0.1) between trait associated targets and phenotypically related disease risk gene sets (**Fig. 4g, Supplementary Table 7**). For example, drug targets enriched among compounds that perturb genes associated with “Hip circumference adjusted BMI” are enriched for risk genes for weight gain, nausea, and psychological stress, and drug targets enriched among compounds that perturb “Coronary Artery Disease” associated genes are enriched for risk genes for heart disease, hypercholesterolemia, abdominal obesity, and myocardial infarction.

Taken together, these chemogenomic enrichments illustrate the potential for the approach described in this study to inform drug discovery and drug development efforts. The identification of side-effect and drug target enrichments linked to known or plausible trait biology supports the veracity of the repurposing predictions, and, more broadly, the power of integrative genomics approaches to identify novel molecular networks that underpin disease.

### Trait-trait correlations and gene-trait associations

To further understand the trait relatedness, we construct a network based on pairwise trait comparison of genetically regulated expression (GReX) (including traits with more than 50 significant associations). By using a broad categorization of traits (**Supplementary Table 4**), we identify 245 pairs of shared gene associations across trait categories and 66 pairs within trait categories (**Fig. 5a, Supplementary Table 8**). Higher numbers of genes are shared between traits that belong to the same trait category than traits belonging to different trait categories; the highest number of genes are shared between low density lipoprotein and total cholesterol in the “lipids” category. Previous studies have shown significant genetic correlation among common traits^39,40^. Pairwise trait GReX correlation shows a positive association with genetic co-heritability^39,40^ (Pearson’s *r* = 0.8, *p* value < 2.79 × 10^-126^) (**Fig. 5b**), extending the genetic similarity among traits to specific genes.

**Figure 5.**
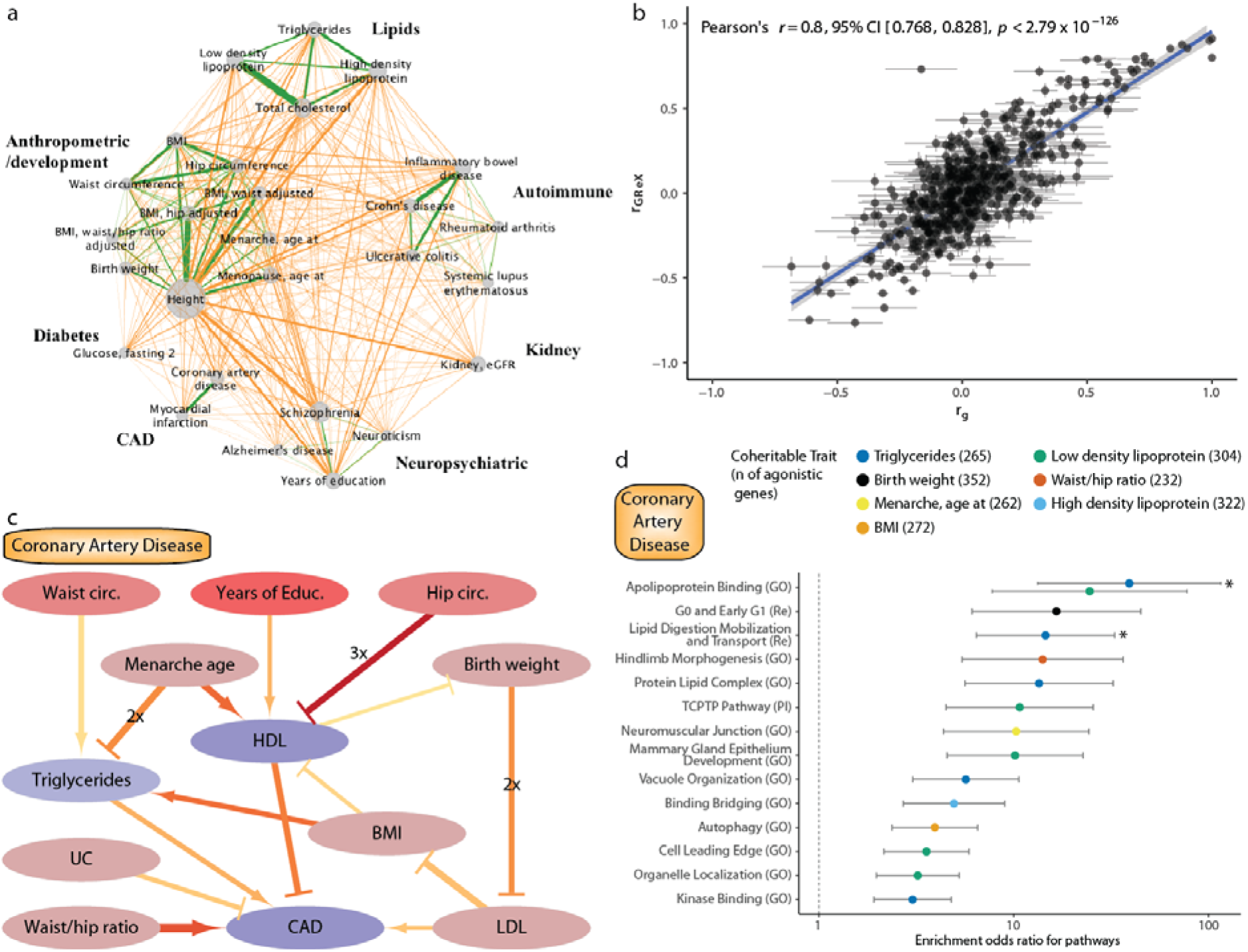
Trait-trait correlations and gene-trait associations. **(a)** Network indicating shared genes within/across trait categories. Only traits that have more than 50 associated genes are showcased. Edge width denotes number of shared genes for each trait pair. The node size indicates number of gene-trait associations for a given trait. Green edges denote within-category trait associations and orange edges denote across-category trait associations. The analysis is based on significantly associated genes with FDR ≤ 1 %. **(b)** Scatter plot of genetic correlation (rg) and genetically regulated gene expression (rGReX) for each pairwise trait combination. Standard error is shown with grey lines, r_g_ and r_GReX_ are highly correlated (Pearson’s r = 0.8, *p* value < 2.79 × 10^-126^). **(c)** Causal trait network of CAD. CAD and up to two traits upstream are plotted in this network graph to demonstrate causal (arrows) and protective (barheaded lines) relationships as estimated by bi-directional regression analysis. The trait nodes are colored based on the parent causal trait network of all the traits of the study (**Supplementary Fig. 16**); nodes that are more times parent and child nodes are a darker shade of red and blue, respectively. In edges, width denotes absolute beta, redder color denotes lower *p* value, and the 2x or 3x labels denote that the relationship is identified in 2 or 3 tissues, respectively. The analysis is based on genes with FDR ≤ 1 %, and only the relationships with p value ≤ 0.05 are shown. **(d)** Graph depicting the odds ratio of pathway enrichment for CAD agonistic genes shared with traits involved in the causal network. Briefly, for causal traits, a list of genes (with unadjusted *p* value ≤ 0.05) that are predicted to change to the same direction (or the opposite direction for protective traits) is used for GSEA for common pathways. In this graph only the top 15 (based on *q* value) results are shown and are ranked based on odds ratio; an asterisk (*) indicates results that have *q* value ≤ 0.05. Error bars represent 95% CI for each enrichment.

We then apply bi-directional regression analyses^41^ on the GReX of different traits across all tissues to infer causal relationships among pairs of traits with significant genetic and GReX correlation (**Fig. 5c** for CAD and **Supplementary Fig. 16** for all the traits in our study). We find evidence that CAD is a complex trait whose predicted gene expression changes can be partly, but directly, explained by predicted expression changes found in individuals with elevated triglycerides, elevated LDL, and increased waist/ hip ratio. On the other hand, predicted expression changes in individuals with increased HDL, or those suffering from ulcerative colitis (UC), are expected to normalize expression changes in individuals with CAD. By expanding the causal network to include more upstream traits, we can see that another 6 traits (waist and hip circumference, years of education, age at menarche, birth weight, and BMI), which are correlated, or anti-correlated, with CAD may cause, or protect, from the predicted expression changes through effects on intermediate traits. For example, waist circumference acts via a causal relationship with triglycerides; other traits follow multiple pathways such as age at menarche, which opposes predicted transcriptomic changes of the increased triglycerides group while promoting imputed transcriptomic changes for individuals with high HDL. We then leverage these causal networks to dissect the pathogenesis of CAD by identifying the molecular pathways shared among all the involved trait pairs. For each trait that can cause or protect from CAD, we identify the “agonistic” genes - genes whose predicted expression is changing towards the same or opposite direction for causal (e.g. triglycerides) and protective (e.g. HDL) traits, respectively. Gene set enrichment analysis of agonistic genes for biological pathways point towards biologically relevant processes for CAD (**Fig. 5d**). For example, a subset of CAD genes (n= 256 out of 2806 genes with *p* value ≤ 0.05) is shared with triglycerides and affects biological processes related to apolipoprotein binding and lipid digestion, mobilization and transport.

Taken together, the pairwise GReX trait correlations illustrate the potential to identify genes that are shared among genetically correlated traits. Agonistic versus antagonistic pleiotropy among two traits can be differentiated by leveraging the directionality of gene expression association in each trait. For traits, such as CAD, this analysis can be applied to dissect the complex phenotype, to identify genes and pathways that are shared with another trait, and potentially identify and develop therapeutic strategies to reverse those perturbations.

## Discussion

The maps of gene expression and regulatory annotations, generated by projects such as REMC^18^, CommonMind^21^, GTEx^23^, and STARNET^22^ hold the potential to further our understanding of non-coding risk genetic variation. Here we describe EpiXcan which, compared to PrediXcan, integrate biologically relevant data in a single framework to improve predictive performance of transcriptome imputation. EpiXcan is also better powered to identify clinically significant results such as enrichment for loss-of-function intolerant genes in neuropsychiatric traits^31,32^ and can detect more robust gene expression changes in genes associated with severe forms of the trait. We apply EpiXcan prediction models from 14 tissues in 58 common and complex traits and examine properties of those associations.

First, gene associations are predominantly identified in pathophysiologically relevant tissues and most associations are only identified in one tissue. Considering that the average correlation between genetically regulated gene expression of unrelated tissues such as blood and brain across 58 traits is 0.38 - 0.42 (Spearman’s *ρ*), we highlight the need for trait-relevant tissue datasets for such studies to be more effective. Second, among genes associated with the traits in this study, we observe significant enrichment for biological pathways involved in trait pathophysiology. Moreover, gene-trait associations are significantly enriched for: (1) pathogenic (or likely pathogenic) genes for the given trait (clinVar), (2) genes associated with trait-relevant phenotypes (SoftPanel), (3) genes that have been associated with clinical signs relevant to the trait (OMIM CS), and (4) ortholog mouse genes with phenotypes that belong to the same phenotypic category as the given trait. This suggests that common variants partly act via smaller effect size perturbations in genes that lead to more severe forms of the phenotype when subject to larger effect size disruptions, as recently similarly suggested^42^.

Third, by leveraging trait-specific transcriptomic changes, we identify known and novel compounds that can reverse trait-specific changes, pointing to potential drug repurposing candidates. To our knowledge, there is only one recent study^43^ that applied a similar approach but it was much more limited in scope (brain tissue – 10 regions - transcriptomic imputation with S-PrediXcan for psychiatric traits) and did not yield any statistically significant results for schizophrenia. In contrast, our study identifies one statistically significant result (phenformin, **Supplementary Table 7**) that is a very potent antidiabetic agent (no longer FDA-approved due to safety concerns) which is not surprising given that glucose homeostasis is altered from illness onset in schizophrenia^44^. Within the top 10 results for schizophrenia we also identify a potent antipsychotic (prochlorperazine), a voltage-gated sodium channel^45^ inhibitor (pramocaine) and guanfacine which was trialed for cognitive impairment in schizophrenia and found to be worthy of further investigation in order to target spatial working memory and continuous performance test reaction time for patients on atypical neuroleptics^46^. It is hard to directly compare the results of the two studies since our approaches differ on many levels: we use EpiXcan, train models on different tissues and more tissue types, and employ a different drug repurposing pipeline. Towards validating our approach, chemogenomic enrichment analysis reveals trait-specific, phenotypically related, side-effects and drug target enrichment for risk associated genes of phenotypically related traits.

Finally, we use bi-directional regression analysis^41^ to construct putative causal trait networks. Causal trait networks built on top of EpiXcan are sufficiently powered to provide valuable insight into the development of complex traits such as CAD. For example, we find that high BMI can influence CAD by two distinct pathways; (a) by positively influencing triglycerides (TG) which would positively influence CAD, and, conversely, (b) by negatively influencing HDL which would negatively influence CAD. The independent effect of BMI on TG and HDL has been shown in a population with a broad spectrum of BMI values^47^ which – as in our study – found no effect of BMI on LDL levels. Downstream, there is genetic evidence to suggest a causal influence of TG on CAD^48^. In addition, a negative correlation of HDL with CAD has been established in observational epidemiology, although a link between genetic loci causal for high levels of HDL and protective for CAD is, at present, elusive^49^. The construction of these causal trait networks allows us to identify genes, among causally-linked traits, that exhibit agonistic pleiotropy participating in shared pathways. Such information could potentially be used to develop distinct therapeutic strategies based on individual comorbidities.

Overall, the described method utilizes epigenomic information to further improve prediction of transcriptomes and it provides a framework for TWASs, improved interrogation of trait-associated biological pathway involvement, and a platform for drug repurposing and treatment development.

To facilitate interpretation, we provide the EpiXcan pipeline, trained models and resulting data tables as an online resource.

## Online methods

### Genotype and expression data

Genotype datasets (CMC, GTEx and STARNET) are uniformly processed for quality control (QC) steps before imputation. We restrict to samples with European ancestry (**Supplementary methods**). Genotypes are imputed using the University of Michigan server^50^ with Haplotype Reference Consortium (HRC) reference panel^51^. RNAseq gene level counts are adjusted for known and hidden confounds, followed by quantile normalization. For CMC gene expression, we use the gene level counts generated from DLPFC RNAseq data^21^ (http://commonmind.org/). For GTEx^52^, we use publicly available quality-controlled gene expression datasets from GTEx consortium (http://www.gtexportal.org/). RNAseq data for STARNET were generated as described in Franzén, *et al*^22^. Additional information for CMC, STARNET and GTEx tissues (for both predictors and observed datasets) including sample sizes is shown in **Supplementary Table 1**. To compare the prediction accuracy of the CMC-trained predictors, we utilize expression data from the HBCC (n = 280 samples^21^) as well as 13 brain areas from GTEx^52^ (**Supplementary Table 1**).

### SNP priors calculation and rescaling to WENet penalty factors

To leverage epigenomic information, we incorporate rescaled SNP priors as penalty factors into a weighted elastic net model. First, we compute eQTLs using MatrixEQTL^53^. Then, epigenome annotations from REMC^18^ are integrated to obtain SNP priors using qtlBHM^20^ (top panel in **Supplementary Fig. 1; Supplementary Materials and Methods; Supplementary Table 9**). Lastly, the SNP priors are rescaled to penalty factors used in WENet by a data-driven rescaling equation. The optimal rescaling equation is approximated by the best performing quadratic Bézier function, providing both the curve of the rescaling function and the minimum value of the penalty factors. Briefly, to determine the best performing rescaling equation, we simulate genotypes (n=500 samples) using HAPGEN2^54^ and haplotypes from the 1000 Genomes Project^55^. For each gene under consideration, we utilize a shifting window policy to generate quadratic Bézier rescaling equations. In each separate window, we define a minimal penalty factor (**Supplementary Fig. 17**) and within that window we evaluate possible intermediate Bézier curve control point locations to test for a wide range of curves for our rescaling equation (**Supplementary Fig. 18**). The equation that exhibits the highest improvement of *R^2^CV* when compared to not assigning penalty factors to the SNPs (as in PrediXcan) is selected. The process to evaluate and select the optimal rescaling equation is described in greater detail in Supplementary Methods.

### Simulation analysis to compare EpiXcan and PrediXcan predictive performance

500 samples are simulated to verify the model performance. For specific gene, suppose ***X*** is the matrix containing genotypes of all cis-SNPs included in the gene. For the *i*-th SNP, we choose an effect estimate **β_*i*_**, so we have vector of estimated effects **β** for all the SNPs of the gene. Gene expression values are simulated by

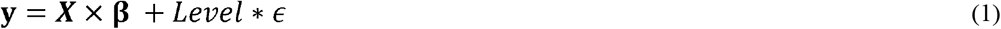

Here ‘×’ denotes matrix-vector product and *ϵ* is normally distributed noise with given standard deviation (SD=0.3). We select ten levels (*Level* from 0.1 to 1) of noise to simulate expression values for given genes. The CMC eQTL beta values are used as the effects in the simulation. We use 1,000 genes with the highest significance from CMC eQTL studies to perform the simulations. For each gene, we simulate 50 times and take the mean value to evaluate the closeness between simulated and real-world gene expressions.

### Large scale gene-trait association analysis

We train predictors of gene expression by applying EpiXcan and PrediXcan to genotype and RNAseq datasets across 14 tissues (**Supplementary Table 1**). For each tissue, we keep genes with pred.perf *q* value of the correlation between cross-validated prediction and observed expression (pred.perf^16^) ≤ 0.01. We identify gene-trait associations, by jointly analyzing summary statistics from 58 complex traits (**Supplementary Table 4**) and gene expression predictors using S-PrediXcan^42^. SNPs in the broad major histocompatibility complex (MHC) region (chromosome 6: 25~35 Mb) are removed. *P* values are adjusted using the Benjamini-Hochberg method of controlling the false discovery rate at ≤ 0.01. The gene-trait associations that remain after this filtering are considered “significant”.

#### Enrichment score

We use an enrichment score^6^ to indicate enrichment or depletion of the trait in a given tissue. For each tissue-trait combination, we count the number of genes that are significantly associated with the trait in that tissue (n_tissue,trait_) and divide them by the number of all the genes that had a prediction either significant or not 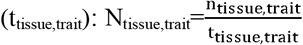. To scale the normalized count N_tissue,trait_ we first subtract the mean normalized count for all tissues for the given trait N_trait_ and then divide the result by the standard deviation (SD) of the normalized count for all tissues for the given trait: 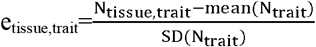. In **Fig. 3b**, e_tissue,trait_ is used as the enrichment score for depicted by the color scale.

#### Unique associations

Uniquely identified genes by EpiXcan (or PrediXcan) are genes that are identified in significant gene-trait associations with one method but not the other. For gene-trait associations found in multiple tissues, we categorize genes as up- (or down-) regulated in the trait if there are more tissues in which the effects are towards the indicated direction. If there are equivalent numbers of tissues in which the gene is positively and negatively correlated with a given trait, we categorize the gene regulation as ambiguous. Transcriptomic imputation yields approximately the same number of genes predicted to be up- or downregulated (z-scores) across each trait (**Supplementary Fig. 19**). To construct the shared gene network in **Fig. 5a**: *(1) we* filter genes so that those with pred.perf *q* values ≤ 0.5% and FDR-adjusted *p* values ≤ 0.5% are retained, *(2)* specifically for shared genes across traits of the same category, we only include genes with high effects (e.g. is |z – score| ≥ mean_*i*_|z – score|_*i*_, *i* number of genes) to limit network density.

#### Novel genes

For identification of novel genes outside of GWAS loci, we define index SNPs based on LD clumped regions using Plink software (v1.9)^56^ The following settings are used: (a) significance threshold for index SNPs is 5 × 10^-8^, (b) significance threshold for clumped SNPs is 5 × 10^-8^, (c) clumping window size is 250 Kb and (d) LD threshold for clumping is 0.1. The coordinates of the GWAS loci are defined as 1 Mb on either side of the index SNP in each clump. The genomic coordinates of the significant genes are then extracted from GENCODE (build GRCh37, release 19) and overlapped with the coordinates of GWAS loci. Those genes that lie outside the overlaps are defined as novel genes. The difference in the number of novel genes identified between the two methods is calculated by subtracting the number of genes identified by PrediXcan from the number of genes identified by EpiXcan. The statistical significance is tested with the null hypothesis that mean difference is not different from zero using one sample t-test (H_0_: *μ*=0).

### Gene set enrichment analyses and phenotypic datasets

To investigate whether the genes associated with a given trait exhibit enrichment for biological pathways, we use gene sets from MsigDB 5.1^57^ and filter out non-protein coding genes as well as genes that do not have eQTL. For the enrichment analysis we only consider traits with >10 genes identified in significant gene-trait associations; this condition is met for 43 traits in our study. Statistical significance is evaluated with one-sided Fisher’s exact test and the adjusted *p* values are obtained by the Benjamini-Hochberg method. Similarly, for **Fig. 2c**, we perform gene set enrichment analysis for all decile bins of pLI from ExAC^26^ (all results can be found in **Supplementary Table 5**). The phenotypic datasets: ClinVar, OMIM CS, SoftPanel, and MGD are prepared as described in **Supplementary Methods** and contain genes that are associated with one or multiple traits. The approximation of known gene-phenotype associations from these datasets allows us to (1) compare the power of EpiXcan vs. PrediXcan in identifying known gene-trait associations (as in **Fig. 2d**) and (2) evaluate the extent to which common risk variants confer trait risk by affecting gene expression levels of genes associated with monogenic forms of the trait or genes associated with similar-to-the-trait phenotypes in humans and mice.

### Computational drug repurposing

We iterated over each trait considered in this study, retaining trait/gene associations with an FDR < 0.1, and converting HGNC gene symbols to NCBI entrez gene identifiers. If a gene is linked with a trait via an association detected in multiple tissues, the associations are summarized as the mean z-score. There are 58 traits with a minimum of 5 positively, and negatively, associated genes and each of these are used as the basis for drug repurposing. For each of these traits, and for each unique compound, we calculate a “connectivity score” based on a modified Kolmogorov-Smirnov score^34^, which summarizes the transcriptional relationship to the trait-associated genes. We estimate statistical significance by generating an empirical Kolmogorov-Smirnov score distribution from the query signature against 1,000 permuted drug signatures. Compound profiles are sourced from Connectivity map^34^. We download and merge the 6,100 individual experiments into a single representative signature for the 1,309 unique small molecule compounds according to the prototype-ranked list method^58^.

#### Chemogenomic enrichment analysis

For each trait, connectivity scores are then used to sort the list of 1,309 compounds and used as the basis for a chemogenomic enrichment analysis. For each compound in the drug signature library, we collect diverse chemogenomic annotations, such as drug target information, side-effect, and therapeutic class associations. Side-effect associations are downloaded from Offsides^59^ and SIDER^59^ and connected to compounds in Connectivity map via Stitch identifiers. Drug target associations include targets referenced in DrugBank^37^, and also an augmented set of associations, based on predictions generated using the Similarity Ensemble Approach^38^. For each of these features, we calculate a signed running sum enrichment score, which reflects whether that feature is over-represented at the extreme ends of the drug list that has been ordered according to trait. Statistical significance of enrichment scores is based on comparison to a large distribution of permuted null scores, generated by calculating scores from randomized chemogenomic sets that contain an equivalent number of compounds to the true set being evaluated. *p* values are adjusted using the Benjamini-Hochberg method of controlling the false discovery rate.

#### Disease risk gene enrichment analysis of trait associated drug targets

We compile disease and trait risk associations from multiple sources, including HGMD^60^, ClinVar^27^, dbGAP^61^, Genetic Associations Disease^62^, GWAS catalog^63^, GWASdb^64^, Human Phenotype Ontology^65^, HuGE^66^, and OMIM^67^. Many of these are accessed through Harmonizome^68^. We use a Fisher’s exact test to compare each set of trait-associated drug targets (that contain at least 3 targets), with each disease risk gene set. The analysis is performed against a background of 2,802 genes, representing the unique set of human drug targets in the combined set of referenced and predicted targets associated with the 1,309 compounds. Two-sided *p* values are adjusted using the Benjamini-Hochberg method of controlling the FDR.

### Trait co-heritability analysis

#### Tissue clustering

To calculate the genetically regulated gene expression correlation (r_GReX_), as shown in **Fig. 3a**, we keep the significant imputed gene expression change (*z* score) values with *q* value ≤ 0.01 and perform pairwise tissue Spearman correlation analysis of the complete cases of *z* scores. To cluster the tissues together for plotting, we use hierarchical agglomerative clustering analysis with Ward’s method.

#### Genetically regulated gene expression correlation (r_GReX_)

Pairwise genetic correlation (r_g_), as shown in **Fig. 5b**, among traits analyzed by GWAS is taken from previously published reports^39,40^. For trait comparisons that appear in both studies we use the more recent study^40^. We consider the genetic correlation between traits significant if *q* value 0.05. To calculate rGReX, we keep the imputed gene expression values with unadjusted *p* value 0.05 and perform pairwise trait Spearman’s correlation analysis with Holm’s adjustment for multiple comparisons. To estimate the correlation of r_g_ and r_GReX_ for the trait pairs in our study we perform Pearson’s correlation analysis with Holm’s adjustment for multiple comparisons.

#### Bi-directional regression and exploratory pathway analyses for putatively causally linked traits

We identify all the significantly correlated trait-pairs (r_g_ and r_GReX_, *q* value ≤ 0.05 as above) and perform bi-directional regression analyses^41^ to identify causal relationships among the traits of our study (**Supplementary Fig. 16**). Then taking as an example the coronary artery disease (CAD), we graph all the putative causal and protective relationships up to 2 nodes upstream in **Fig. 5c** (when the causal relationship is bi-directional between 2 traits, the relationship with the higher degrees of freedom is kept) and perform pathway enrichment analysis of shared agonistic genes for this causal network in **Fig. 5d**. For each causal or protective trait in the network, we generate a list of genes whose expression changes are predicted towards the same direction (or the opposite direction for protective traits) in CAD. These lists of shared “agonistic” genes are used for GSEA for common pathways. In **Fig. 5d**. only the top 15 (based on *q* value) results are shown and are ranked based on odds ratio.

## URLs

CMC: http://commonmind.org/

Synapse for CMC data: https://www.synapse.org/cmc

GTEx portal: http://www.gtexportal.org/

MSigDB: http://software.broadinstitute.org/gsea/msigdb

EpiXcan website and repository: http://icahn.mssm.edu/EpiXcan

EpiXcan source code: https://bitbucket.org/roussoslab/epixcan

qtlBHM package: https://github.com/rajanil/qtlBHM

RHOGE package: https://github.com/bogdanlab/RHOGE

PrediXcan pipeline: https://github.com/hakyim/PrediXcan

PredictDB resource: https://github.com/hakvimlab/PredictDB_Pipeline_GTEx_v7

## Acknowledgments

Authors would like to thank Scott Dickinson, Alvaro Barbeira, and Hae Kyung Im for providing valuable feedback on their PrediXcan and S-PrediXcan pipelines.

This work was supported by the National Institutes of Health (NIH) grants R01AG050986 (P.R.), R01MH109897 (P.R. and E.E.S.) and R01MH109677 (P.R.), and the Veterans Affairs Merit grants BX002395 and BX004189 (P.R.).

## Author contributions

P.R., W.Z. and G.V. conceived and designed the study. W.Z. implemented the EpiXcan pipeline and built the website database. W.Z., G.V., V.R. and B.R. did the downstream analysis. Additional feedback and discussion was provided by J.T.D., E.E.S., J.L.M.B., Y.K., J.F.F., G.E.H. W.Z., G.V., V.R. and P.R. wrote the manuscript with input from all the authors.

