## Supplementary text and figures for "Integrative transcriptome imputation reveals tissue-specific and shared biological mechanisms mediating susceptibility to complex traits"

**This PDF file includes:**

Materials and Methods

Supplementary Text

Theorem and proof

Supplementary Figs. 1 to 23

Supplementary Tables 1 to 10

### Materials and Methods

#### Weighted elastic net (WENet) model utilized in EpiXcan

The elastic net (ENet) linear regression model is implemented in PrediXcan[1]. Criterion can be written as (equation (S.1))

$$\mathbb{C}_{\text{ENet}}(\boldsymbol{\theta}, \lambda, \alpha) = \sum_{i=1}^{n}[y_i - X_i\boldsymbol{\theta}]^2 + \lambda\alpha|\boldsymbol{\theta}|_1 + \lambda(1-\alpha)|\boldsymbol{\theta}|_2\,, \tag{S.1}$$

where $X_i$, $1 \leq i \leq n$, is the $i$-th row-vector of matrix $\boldsymbol{X}$ containing genotypes with dosages from 0 to 2. $n$ is the number of samples. In (S.1) all SNPs are equally treated. In EpiXcan, we use a weighted ENet (WENet) model that incorporates penalty factors from rescaled SNP priors, the criterion of which can be written as (equation (S.2)):

$$\mathbb{C}_{\text{WENet}}(\boldsymbol{\theta}, \lambda, \alpha) = \sum_{i=1}^{n}[y_i - X_i\boldsymbol{\theta}]^2 + \lambda\alpha|\boldsymbol{\theta}|_{\boldsymbol{w}} + \lambda(1-\alpha)\boldsymbol{\theta}^T\mathbf{W}\boldsymbol{\theta} \tag{S.2}$$

In (S.2), $\mathbf{W}$ is the weight matrix that stores the penalty factors for SNPs. $|\boldsymbol{\theta}|_{\boldsymbol{w}} = \sum_{j=1}^{m} w_j|\theta_j|$, with $w_j$ corresponding to the penalty factor of the $j$-th SNP. $m$ is the number of *cis*-SNPs.

In equation (S.1), $n$-by-$m$ matrix $\boldsymbol{X}$ encloses genotype of *cis*-SNPs of the specific gene for all samples, i.e., there are $n$ samples and $m$ *cis*-SNPs. $|\boldsymbol{\theta}|_1$ is the $L_1$ norm of $\boldsymbol{\theta}$, which is coefficient vector of SNPs. $\boldsymbol{y}$ contains expression values of the specific gene for all the samples. $y_i$ is the $i$-th entry of the response vector $\boldsymbol{y}$, which includes the expression value of the gene for the $i$-th sample. For later presentations, we use $\mathbf{x}_{\boldsymbol{j}}$, $1 \leq j \leq m$, to denote the $j$-th column vector of $\boldsymbol{X}$. Following the definitions, we know $\mathbf{x}_{\boldsymbol{j}}$ encloses the genotype of the $j$-th SNP with respect to all the samples. The $\alpha$ parameter is set to 0.5 and $\lambda$ is estimated via cross-validation (CV).

In equation (S.2), $\mathbf{W}$ is a diagonal matrix and its entries are the penalty factors that utilized. We see from equation (S.2) that if $\mathbf{W} = \mathbf{I}$, which is identity matrix, the algorithm becomes the standard traditional elastic net module (S.1). From this perspective, the WENet model is more general and it consists of classic ENet as a special case. Equation (S.2) contains three terms: the negative log-likelihood function of linear regression; the $L_1$ normalized term, which penalizes the $L_1$ norm of $\boldsymbol{\theta}$; and the ridge penalty, which can be formulated as the inner product of $\boldsymbol{\theta}$ with respect to matrix $\mathbf{W}$, $<\boldsymbol{\theta}, \boldsymbol{\theta}>_{\mathbf{W}}$. In case when $\alpha = 1$ and $\lambda \neq 0$, the method is reduced to Lasso. If $\lambda = 0$, the method is even more simplified as a standard regression model without penalties. If all the penalty factors are 1's, *i.e.*, matrix $\mathbf{W}$ is identity matrix, the model is reduced to standard ENet without penalty weights.

The model employed by the elastic net method in Gamazon *et al*[1] is based on the criterion (S.1), where all SNPs have the same penalty factor, which is set to 1 by default. Grouping effects[2,3] of WENet model are provided in **Theorem and proof** (Theorem 1).

#### SNP priors

We first prepare eQTL statistics (computed with MatrixEQTL[4]) and SNP annotations (extracted from REMC https://egg2.wustl.edu/roadmap/web_portal/). For each eQTL tissue, we use the matched REMC tissue to extract the corresponding annotations (**Supplementary Table 9**). We then provide them as input of qtlBHM[6] that utilizes a Bayesian hierarchical model to calculate priors, which is a measure of SNP causality. Priors are derived from chromHMM[5] and for each tissue, SNPs in the same state are assigned the equivalent priors based on the chromHMM tracks that they located in. The REMC tissues that match eQTL tissues and prior statistics for all tissues of this study for each annotation category are provided in **Supplementary Table 9**. SNP priors for a given dataset can be calculated using our pipeline at https://bitbucket.org/roussoslab/epixcan; for the SNP priors included in this study we offer them in the predictor databases as a direct download at https://icahn.mssm.edu/EpiXcan.

#### Data-driven equation that rescales SNP priors to penalty factors

The higher the estimated SNP priors given by qtlBHM[6] the higher the likelihood that the SNP has an important effect on gene expression. On the other hand, higher penalty factors in the WENet model denote a smaller effect in gene expression. Thus, optimal equations must be found to properly rescale priors to penalty factors. For this study, we developed a method based on Bézier curves employing a shifting-window strategy to approximate the data-driven rescaling function. We theoretically can have a different rescaling equation for each gene but for this study we opt to use one rescaling equation for all models of a given tissue for simplicity and computational resource efficiency. The steps of this method using the CMC tissue dataset as a template are as follows:

1) We perform PrediXcan and obtain the target $R^2_{CV}$ for all genes. We then select 8 genes that are representative for different levels of $R^2_{CV}$ that can be found in the study. For CMC we select the following genes:

| Gene symbol | Target $R^2_{CV}$ |
|---|---|
| ***DDX11*** | 0.7631 |
| ***ADAM15*** | 0.3029 |
| ***C1orf112*** | 0.1497 |
| ***C1RL*** | 0.5098 |

| | |
|---|---|
| ***ERBB3*** | 0.0204 |
| ***ECT2L*** | 0.0131 |
| ***SEPT1*** | 0.0053 |
| ***ZNF346*** | 0.0050 |

2) We simulate 500 genotypes using HAPGEN2[7]. We use haplotypes from the 1000 Genomes Project[8] and fine-scale recombination map[7] to simulate genotypes. We further filter the genotypes to include SNPs with MAF of at least 5%. We then keep the SNP structure for the *cis* SNPs of the 8 genes that selected.
3) For each of the genes from (1), we perform simulations to select best rescaling function. All rescaling's are based on quadratic Bézier curves. A *n*-th order Bézier curve is defined by a set of points, $P_0$ through $P_n$, which are control points. The first ($P_0$) and last ($P_n$) control points are usually called the starting and ending control points of the curve. All other points are intermediate control points that do not lie on the curve, however they decide the shapes of the curve. A *n*-th order Bézier curve has *n*+1 control points, *n*-1 of which are intermediate control points - a brief introduction of Bézier curves is enclosed in the next section. We then:
    a. Define region of prior-to-penalty factor mapping. As shown in **Supplementary Fig. 17b**, to map priors to penalty factors, we first define an area with x $\in$ [0, maximal SNP prior] corresponding to the SNP priors and y $\in$ [0, 1] corresponding to penalty factors.
    b. Shifting-window policy. We then divide the rectangle region of prior-to-penalty factor mapping into several sub-windows. In each of the sub-windows, we have ranges of both the penalty factors and the priors. As described above, penalty factors should decrease with increasing values of priors so that important SNPs can have a larger effect on transcriptomic imputation. We set the upper bound of penalty factors at 1 (as in the ENet model employed by PrediXcan) to which the minimal value of priors will be mapped (**Supplementary Fig. 17b**). Since we do not have a lower bound of the penalty factors to which the maximal priors will be mapped, we map the maximal prior to the lowest rescaled factor value ($y_2$) in each sub-window starting from 0 and going all the way to 1 with step size 0.1 (step size is arbitrarily set) (e.g. in **Supplementary Fig. 17a**, $y_2$=0.5).
    c. Define a set of possible rescaling equations in each sub-window. Above, we set $P_0$ (0, 1), the starting point of the curve, and $P_2$ ($x_2$, $y_2$) where $x_2$ is the maximal point of the priors and $y_2$ with range [0, 1] that is fixed for each sub-window (**Supplementary Fig. 17**). For each sub-window we define a grid (we set grid size of 0.1), denoting all the possible positions for $P_1$ intermediate control points that we will evaluate. For each different intermediate control point ($P_1$), we get a quadratic Bézier rescaling equation. An example is illustrated in **Supplementary Fig. 18** (only a

limited number of candidate rescaling functions are shown, although there are hundreds of possibilities).

d. Perform simulations to assess performance for each rescaling equation. We apply all possible Bézier curves as rescaling candidates to compute the $R^2_{CV(simulation)}$ for 100 times of gene-specific simulations. For each simulation we use the 500 simulated genotypes from (2) and assess performance ($R^2_{CV(simulation)}$) against simulated gene expression. Simulated gene expression is calculated by equation (2). Instead, we choose PrediXcan predictors as the effect estimates, and *noise* is normally distributed with given standard deviation (SD=0.3).

e. Select the best performing rescaling equation in each sub-window from (3b). The rescaling function with highest improvement of $R^2_{CV(simulation)}$ from (3d) is chosen as the optimal rescaling in the sub-window. We select for maximal improvement as determined by:

$$\Delta R^2_{CV} = mean(R^2_{CV(simulation)} \text{ of EpiXcan}) - mean(R^2_{CV(simulation)} \text{ of PrediXcan}) \quad \text{(S.3)}$$

In **Supplementary Fig. 20**, we show the optimal rescaling equations for each sub-window for gene *DDX11*.

f. Select the best sub-window-specific performing rescaling equation from (3e) for each gene (from (1)). For every optimal rescaling equation from each sub-window (e.g. **Supplementary Fig. 20**), we perform another 100 simulations as described in (3d). And select the best performing one based on equation (S.3), which is the optimal rescaling equation for the gene.

g. Select the best gene-specific performing rescaling equation to use for the tissue. For each of the 8 genes from (1), we can see the best performing rescaling equations (**Supplementary Fig. 21**). We perform EpiXcan using each one of these rescaling equations and select the best performing one based on $R^2_{CV}$ improvement.

Finally, to conserve computational resources, we skew the rescaling equation from CMC, to fit the maximal priors (**Supplementary Table 9**) for all other tissues (**Supplementary Fig. 22**). Here, we provide a framework for data-driven adaptive rescaling. Depending on the needs of each study, researchers may opt to estimate from scratch tissue-specific rescaling equations or even use gene-specific rescaling equations.

#### Bézier curves with interpolation functions

The $n$-th order Bézier curve is determined by $n$+1 control points, of which $n$-1 are intermediate control points. For briefness, we list quadratic Bézier curves here and the function is given as

$$y = (1-t)^2 y_0 + 2t(1-t)y_1 + t^2 y_2, \quad x = (1-t)^2 x_0 + 2t(1-t)x_1 + t^2 x_2 \quad \text{(S.4)}$$

As variable $t$ varies from 0 to 1, $y$ is a function of $x$ with second order. $t$ here is an intermediate variable, which is used to define Bézier function. $x_k$ and $y_k$, $k$=0,1,…,$n$ are the coordinates of control points.

Note that in equation (S.4), $x$ and $y$ are denoted by separate functions with respect to variable $t$. We deduce a function of $y$ with respect to $x$ by eliminating $t$ to calculate rescaled values, which are the penalty weights or factors that used in EpiXcan. Here $x$ stores primitive priors, from which we obtain the penalty factors $y$.

As we stated earlier, more intermediate control points are necessary for higher order Bézier interpolations. Theoretically, the more control points we use and the higher order of the interpolations, the higher accuracy will be achieved. To balance accuracy and computation complexity, we use quadratic (second order/degree) Bézier interpolations to approximate the rescaling equations and employ them to the EpiXcan approach. Selecting quadratic Bézier interpolation also has the benefit of not having to control for counter-intuitive increase of penalty factors with increase in priors that can happen in the middle of the curve with higher order equations.

#### Genotype preprocessing

We remove samples with call rate < 0.95, sex mismatch (genetic sex different from pedigree sex) and autosomal heterozygosity deviation (|Fhet|>0.2). We remove variants with call rate < 0.95, Hardy-Weinberg equilibrium (HWE) P < $1.0\times10^{-6}$. We identified related individuals using identity by descent analysis (IBD). One of each pair of related individuals (piHAT > 0.2) is removed at random. Since the entire STARNET cohort and the majority of the CMC cohort include individuals of EUR ancestry, we use genotype and gene expression information only from this population to train the models. For this, we merge the samples with 1000 genome EUR subset and do principal component analysis (PCA) using ~25,000 pruned and thinned variants. We plot first and second principal components and define an ellipsoid based on 1000G EUR samples (**Supplementary Fig. 23**). Those that lie 8 SD away from the center of this ellipsoid are considered as genetic outliers and removed (**Supplementary Fig. 23**). For post imputation, we remove variants with INFO < 0.8, minor allele frequency (MAF) < 0.01, more than one alternative allele. We also remove ambiguous alleles (A/T, G/C), indels (insertions and deletions) and variants without RS identifier.

#### Estimating adjusted $R^2$

We compare the performance of EpiXcan and PrediXcan models using adjusted $R^2$ (for both $R^2_{\mathrm{CV}}$ and $R^2_{\mathrm{PP}}$), which control for sample size of the training dataset (same for both methods) and for the number of predictors in the model (differs for each gene between methods). We group the training samples *a priori* prior the cross validation and use the same groupings in both EpiXcan and PrediXcan. The adjusted $R^2$ is computed using formula (S.5) where $R^2$ is either $R^2_{CV}$ or $R^2_{PP}$, $n_{\mathrm{sample}}$ is the sample size and $n_{\mathrm{SNP}}$ is number of SNPs in the model. For correlation $R^2_{PP}$ adjustments, we use the $n_{\text{sample}}$ and $n_{\text{SNP}}$ in the source models (predictors, **Supplementary Table 1**).

$$R^2_{\text{adj}} = 1 - \frac{(1-R^2)(n_{\text{sample}}-1)}{n_{\text{sample}}-n_{\text{SNP}}-1} \quad \text{(S.5)}$$

We use Wilcoxon and one-sample sign tests for the statistical comparisons of adjusted $R^2_{\text{CV}}$ between PrediXcan and EpiXcan models.

#### Enrichment of clinically significant genes

To compare the clinical significance of the gene-trait associations identified by EpiXcan and PrediXcan, we compile sets of known gene-trait associations by utilizing five different archives:

1. ClinVar[9]: a public archive of relationships among human sequence variation and phenotypes. We only keep the subset of entries that: a) have at least one current submission interpreting as pathogenic or likely pathogenic, b) provide a gene name, c) provide a phenotype. This dataset allows trait-specific gene associations that have high confidence but returns a limited number of genes for each trait.

2. OMIM CS (OMIM Clinical Synopses): a custom subset of the OMIM[10] compendium. This subset is constructed by keeping the genes from the above clinVar dataset that have a corresponding OMIM ID associated with the entry. Then, by using the OMIM API, the Clinical Synopsis Data are fetched for each OMIM ID to allow us to query trait-specific association of relevant clinical signs. Genes are thus linked with clinical signs from a big subset of genetic disorders allowing for a greater number of gene-trait associations when compared with ClinVar.

3. SoftPanel[11]: a method for grouping diseases and related disorders for generation of customized diagnostic gene panels. For traits that have a corresponding ICD-10 number, we use the respective disorder or disorder group and extract the relevant gene sets. For traits that the latter extraction method does not yield any genes (either due to no ICD-10 classification equivalent or due to lack of genes identified with that method), we use the keyword-based search of SoftPanel which queries the OMIM database for keyword-matching disorders. The underlying design of the tool allows for even "softer" associations of the genes with the trait thus providing a larger trait-specific list of genes when compared with the clinVar dataset and OMIM CS.

4. MGD (MGI Phenotypes)[12]: this dataset contains gene-phenotype associations from mouse lines. We can thus infer trait-specific gene associations for the respective human ortholog genes. Direct phenotype overlap with human traits is challenging, in most cases the mouse phenotype is more descriptive and does not use names of human diseases, disorders or syndromes; therefore, phenotype categories are used to query this database.

5. pLI (by ExAC)[13]: this dataset provides probabilities of loss of function intolerance (pLI) for each gene; the higher the pLI the higher the likelihood that this gene performs an essential function. This dataset does not provide trait-specific information but serves as an unbiased dataset to rank the "indispensability" of the genes.

#### Preparation of the datasets

**ClinVar (dataset 1)**. The goal is to prepare a table that lists genes in clinVar that are likely pathogenic or pathogenic and associate them with traits; the following process was performed in January 2018: (1) the clinVar variant summary tabular file was downloaded from NCBI, (ftp://ftp.ncbi.nlm.nih.gov/pub/clinvar/tab_delimited/variant_summary. txt. gz), (2) arrays of genes are excluded, (3) variants without gene names are excluded, (4) variants without associated phenotypes are excluded, (5) variants that don't have at least one current submission interpreting it as likely pathogenic or pathogenic are excluded, (6) the table is aggregated at the gene and phenotype level. The column "PhenotypeList" provides the phenotypes that are used for association with traits of our study, queries are performed as described in **Supplementary Table 10**. 135 unique genes from EpiXcan predictions are directly associated with our traits based on the query table below:

**OMIM CS** (**dataset 2**). Briefly, the clinVar dataset (**dataset 1**) was used as a scaffold and it was populated with information of clinical synopses from OMIM in January 2018 as follows: (1) only the genes that have an associated OMIM ID were kept, (2) we obtained an OMIM API key and did API calls to receive clinical synopsis information for each OMIM ID while respecting call limitations to reduce server load (https://omim.org/help.api) by enforcing a sensible in-between calls time delay, (3) the acquired data were used to populate the table with clinical signs information, (4) only the genes that have OMIM_CS (clinical synopsis) information are queried. Depending on the trait, specific keywords are used to search within the clinical synopsis data (**Supplementary Table 10**) and the identified genes are associated with the trait. 542 unique genes from EpiXcan predictions are associated with our traits.

**SoftPanel (dataset 3).** SoftPanel[11] is an online tool that generates panels of relevant genes based on several query types such as ICD-10 codes and keyword searches (also utilizing the OMIM API) for diseases and phenotypes. The tool was accessed at http://www.isb.pku.edu.cn/softpanel/ in February 2018. The search terms for the 58 traits in our study are listed in **Supplementary Table 10**. 1,362 unique genes from EpiXcan predictions are associated with our traits.

**MGD (**MGI Phenotypes, **dataset 4**, accessed in June 2018**)**. The MGI phenotypes dataset can be generated with the following method: (1) retrieve the .bb (big bed) files from http://www.informatics.jax.org/downloads/TrackHubs/mm10/ that have phenotype information, (2) convert .bb files to .bed files using the BigBedTo Bed binary (http://hgdownload.cse.ucsc.edu/admin/exe/linux.x86_64/bigBedToBed), (3) use the MGI_IDs from the bed file to query the list of all mouse phenotypic alleles (http://www.informatics.jax.org/downloads/reports/MGI_PhenotypicAllele.rpt) to get the respective MGI Marker Accession IDs, (4) use the MGI Marker Accession IDs to retrieve the (human) ENSEMBL IDs for each gene from a conversion table. (It can be generated at https:// www. genenames. org/cgi-bin/download if the "Mouse Genome Database ID (supplied by MGI)" is included. 1,673 unique genes from EpiXcan predictions are associated with our traits.

**pLI** (Probability of loss-of-function intolerance**, dataset 5).** The generation of this dataset is previously described[13] and the table was downloaded from (ftp://ftp.broadinstitute.org/pub/ExAC_release/release0.3.1/functional_gene_constraint/fordist_cleaned_exac_r03_march16_z_pli_rec_null_data.txt). Data are binned at 10% increments or thresholds as described. By convention we refer to genes belonging to the highest decile as extreme loss-of-function intolerant.

#### Gene set enrichment analysis for pLI

GSEA is performed for all pLI (probability of loss-of-function intolerant, **dataset 5**) deciles, $p$ values are calculated with the fisher exact test and are FDR-adjusted to $q$ values. We first perform GSEA for all the significant genes and then we perform a second separate GSEA for significant genes distributed in 8 lists, one for each trait category. No other pLI decile bins yield statistically significant results ($q$ value < 0.05) as shown in **Supplementary Table 5**.

##### Z-score differences for clinical datasets

The Δ[z] (EpiXcan – PrediXcan) values for all the gene-trait associations that are significant from either EpiXcan or PrediXcan are considered. Each of the 5 panels corresponds to a different clinically relevant dataset (**datasets 1-5**). Of note is that the high pLI subset corresponds to pLI ≥ 0.9 (extreme loss of function intolerant genes) and is by design non-trait-specific (thus the higher number of observations in **Fig. 2d**). *P* value is calculated with the one sample sign test against a theoretical median of 0 ($H_0$: $\tilde{X} = 0$). The ratio is the number of Δ[z] measurements in favor of EpiXcan to the respective number for PrediXcan.

#### GWAS statistics

We download 58 GWAS summaries from public datasets, and the traits are categorized into broad overall categories (**Supplementary Table 4**). If there are multiple versions that are available, we keep the study with the largest sample size in our investigations. For some of the data, such as systemic lupus erythematous, we applied for access to the summary statistics from authors. The statistics for Alzheimer's disease are obtained from the International Genomics of Alzheimer's Project (IGAP), which is a two-stage study based on GWASs of European ancestry. IGAP uses genotyped and imputed data on ~7 million SNPs in stage I to analyze published GWAS datasets consisting of more than 17 thousand Alzheimer disease subjects and 37,154 controls. For the detailed information regarding resources of all the GWASs, please refer to **Supplementary Table 4**.

### Supplementary Text

#### EpiXcan has better performance than PrediXcan

*(1)* Performance evaluation in brain tissue. Dorsolateral pre-frontal cortex (DLPFC) gene expression and CommonMind Consortium (CMC) genotype data are utilized as one of the training sets for our approach. Human brain collection core (HBCC) and Genotype-Tissue Expression (GTEx) brain tissue transcriptome data are used only for verification as test datasets (**Supplementary Table 1**). *(2)* Performance evaluation in cardiometabolic tissues. The Stockholm-Tartu Atherosclerosis Reverse Network Engineering Task (STARNET) dataset for seven tissues and the GTEx dataset for six tissues (same tissues as STARNET excluding mammary artery) serve as training and test datasets, respectively and vice versa (**Supplementary Table 1**).

First, we use cross-validation to evaluate prediction performance. The majority of the reference panel genes (>90%) are contained in the EpiXcan-trained predictor database and the overall $R^2_{CV}$ is better than PrediXcan trained PredictDB's (**Fig. 1, Supplementary Fig. 3**) with significant pair-wise Wilcoxon test *P* value regarding all the datasets that we utilized (**Supplementary Table 2**). We list the numbers of genes with $R^2_{CV} \geq 0.01$ from both models. Using 0.01 as the $R^2_{CV}$ cut-off, we detect more genes with EpiXcan having good performance. In addition, EpiXcan has lower root-mean-square error (RMSE) values, further indicating increased performance (**Supplementary Table 2**).

Finally, we use independent test datasets to evaluate prediction performance and external model validity. We use the CMC dataset[14] to train the brain tissue model using both EpiXcan and PrediXcan. Afterwards, we first use the trained database of predictors (PredictDBs) to predict transcriptomes using HBCC genotype data[14]. We show that when using EpiXcan, higher correlations between predicted and observed expression (in HBCC brain) are obtained (**Fig. 1, Supplementary Fig. 4**), with pairwise Wilcoxon test *P* value $< 9.0 \times 10^{-16}$ (**Supplementary Table 3**). We then use the CMC-trained PredictDB's to predict GTEx brain tissue expression and we compare it with observed expression values from 13 different brain regions in that cohort. For all the brain regions, EpiXcan improves prediction performance (**Fig. 1, Fig. 3, Supplementary Fig. 4**). Similarly, for cardiometabolic tissues, we use 7 trained STARNET models to predict corresponding 6 GTEx transcriptomes and vice versa and, overall, we observe better predictive correlation for EpiXcan-trained models (**Fig. 1, Supplementary Fig. 4, Supplementary Fig. 4**).

#### Theorem and proof

Optimal coefficients in equation (S.2) are estimated by equation (S.6):

$$\widehat{\boldsymbol{\theta}} = \underset{\boldsymbol{\theta}}{\operatorname{argmin}}\, \mathbb{C}_{\mathrm{WENet}}(\boldsymbol{\theta}, \lambda, \alpha) \tag{S.6}$$

Grouping effect of the WENet model is given in Theorem 1.

**Theorem 1**. Suppose $\widehat{\boldsymbol{\theta}} = \underset{\boldsymbol{\theta}}{\operatorname{argmin}}\, \mathbb{C}_{\mathrm{WENet}}(\boldsymbol{\theta}, \lambda, \alpha)$, given data $(\mathbf{y}, \mathbf{X})$ where $\mathbf{X}$ is standardized, and parameters $(\lambda, \alpha)$, if $\widehat{\boldsymbol{\theta}}_i \widehat{\boldsymbol{\theta}}_j > 0$, define $D_{\lambda,\alpha}(i,j) = \frac{1}{|\mathbf{y}|_1} |w_i \widehat{\boldsymbol{\theta}}_i(\lambda,\alpha) - w_j \widehat{\boldsymbol{\theta}}_j(\lambda,\alpha)|$, then $D_{\lambda,\alpha}(i,j) \le \frac{\sqrt{2(1-\sigma)}}{\lambda(1-\alpha)} + \frac{\alpha|w_i - w_j|}{2(1-\alpha)|y|_1}$ .

Here $\sigma$ is sample correlation of $\mathbf{x}_i$ and $\mathbf{x}_j$.

##### Proof

Since $\widehat{\boldsymbol{\theta}}_i(\lambda,\alpha)\widehat{\boldsymbol{\theta}}_j(\lambda,\alpha) > 0$, $\operatorname{sign}(\widehat{\boldsymbol{\theta}}_i) = \operatorname{sign}(\widehat{\boldsymbol{\theta}}_j)$. Because $\widehat{\boldsymbol{\theta}} = \underset{\boldsymbol{\theta}}{\operatorname{argmin}}\, \mathbb{C}(\boldsymbol{\theta}, \lambda, \alpha)$, $\widehat{\boldsymbol{\theta}}$ satisfies $\left.\frac{\partial \mathbb{C}}{\partial \boldsymbol{\theta}_k}\right|_{\boldsymbol{\theta}=\widehat{\boldsymbol{\theta}}} = \mathbf{0}$ if $\widehat{\boldsymbol{\theta}}_k(\lambda,\alpha) \ne 0$. Thus

$$\mathbf{2}(\mathbf{y} - \mathbf{X}\widehat{\boldsymbol{\theta}})^{\mathbf{T}}\, \mathbf{x}_{\boldsymbol{k}} + \lambda\alpha \operatorname{sign}(\widehat{\boldsymbol{\theta}}_k) w_k + \mathbf{2}\lambda(1-\alpha)\widehat{\boldsymbol{\theta}}^{\mathbf{T}}\, W_k = 0$$

Here $W_k$ is the $k$-th column vector of matrix $\mathbf{W}$. Hence

$$\mathbf{2}\left(\mathbf{y} - \mathbf{X}\widehat{\boldsymbol{\theta}}\right)^{\mathbf{T}} \mathbf{x}_{\boldsymbol{i}} + \lambda\alpha \operatorname{sign}\left(\widehat{\boldsymbol{\theta}}_i\right) w_i + \mathbf{2}\lambda(1-\alpha)\widehat{\boldsymbol{\theta}}^{\mathbf{T}}\, W_i = 0 \tag{S.7}$$

$$\mathbf{2}\left(\mathbf{y} - \mathbf{X}\widehat{\boldsymbol{\theta}}\right)^{\mathbf{T}} \mathbf{x}_{\boldsymbol{j}} + \lambda\alpha \operatorname{sign}\left(\widehat{\boldsymbol{\theta}}_j\right) w_j + \mathbf{2}\lambda(1-\alpha)\widehat{\boldsymbol{\theta}}^{\mathbf{T}}\, W_j = 0 \tag{S.8}$$

Subtracting (S.8) from (S.7), we have

$$\mathbf{2}\left(\mathbf{y} - \mathbf{X}\widehat{\boldsymbol{\theta}}\right)^{\mathbf{T}} (\mathbf{x}_{\boldsymbol{i}} - \mathbf{x}_{\boldsymbol{j}}) + \lambda\alpha \operatorname{sign}\left(\widehat{\boldsymbol{\theta}}_i\right)(w_i - w_j) + \mathbf{2}\lambda(1-\alpha)\widehat{\boldsymbol{\theta}}^{\mathbf{T}}\, (W_i - W_j) = 0 \tag{S.9}$$

According to property of matrix **W**,

$$\widehat{\boldsymbol{\theta}}^{\mathbf{T}}\, (W_i - W_j) = \boldsymbol{w_i}\widehat{\boldsymbol{\theta}}_{\boldsymbol{i}} - \boldsymbol{w_j}\widehat{\boldsymbol{\theta}}_{\boldsymbol{j}} \tag{S.10}$$

From (S.9), (S.10) and Cauchy-Schwartz inequality as well as property of $L_1$ norm, we get

$$\left|\boldsymbol{w_i}\widehat{\boldsymbol{\theta}}_{\boldsymbol{i}} - \boldsymbol{w_j}\widehat{\boldsymbol{\theta}}_{\boldsymbol{j}}\right| \le \frac{\mathbf{1}}{\lambda(1-\alpha)} \left|\mathbf{y} - \mathbf{X}\widehat{\boldsymbol{\theta}}\right|_{\mathbf{1}} \left|\mathbf{x}_{\boldsymbol{i}} - \mathbf{x}_{\boldsymbol{j}}\right|_{\mathbf{1}} + \frac{\alpha|w_i - w_j|}{\mathbf{2}(1-\alpha)} \tag{S.11}$$

From Zou et al.[15], we know

$$\frac{\mathbf{1}}{\lambda(1-\alpha)|y|_1} \left|\mathbf{y} - \mathbf{X}\widehat{\boldsymbol{\theta}}\right|_{\mathbf{1}} \left|\mathbf{x}_{\boldsymbol{i}} - \mathbf{x}_{\boldsymbol{j}}\right|_{\mathbf{1}} \le \frac{\sqrt{2(1-\sigma)}}{\lambda(1-\alpha)} \tag{S.12}$$

Both sides of (S.11) being divided by $|y|_1$ and from (S.12) we get

$$\frac{1}{|\mathbf{y}|_1} |w_i \widehat{\boldsymbol{\theta}}_i(\lambda,\alpha) - w_j \widehat{\boldsymbol{\theta}}_j(\lambda,\alpha)| \le \frac{\sqrt{2(1-\sigma)}}{\lambda(1-\alpha)} + \frac{\alpha|w_i - w_j|}{\mathbf{2}(1-\alpha)|\mathbf{y}|_1} \tag{S.13}$$

### Supplementary figures

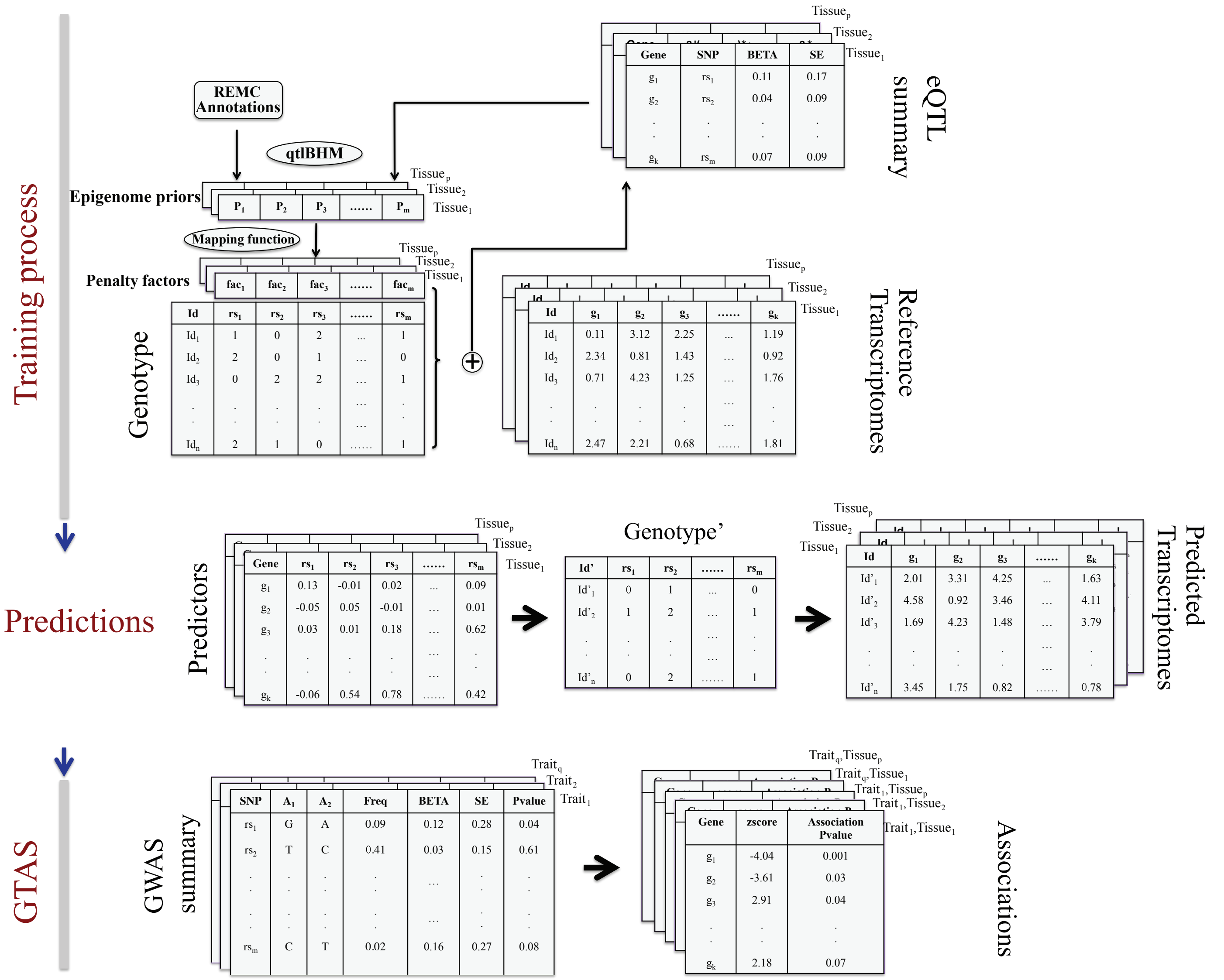

**Supplementary Figure 1. Schematic of the EpiXcan workflow.** For training of the prediction model (top panel), *m* genotypes and *k* transcripts are considered across *n* individuals in *p* tissue datasets. We obtain SNP priors by using a hierarchical Bayesian model (qtlBHM) that jointly analyzes REMC epigenome annotations and eQTL statistics. The priors are then transformed with an adaptive mapping function to penalty factors, which are then utilized by the WENet model. Using the WENet model, we jointly analyze SNP priors, genotypes and gene expression traits to estimate genetically regulated expression component across different tissues. For the gene-trait association studies (bottom panel), we integrate the SNP-transcriptome effect sizes with complex traits effect sizes to estimate the association between predicted gene expression and a trait while taking in consideration the linkage disequilibrium among SNPs.

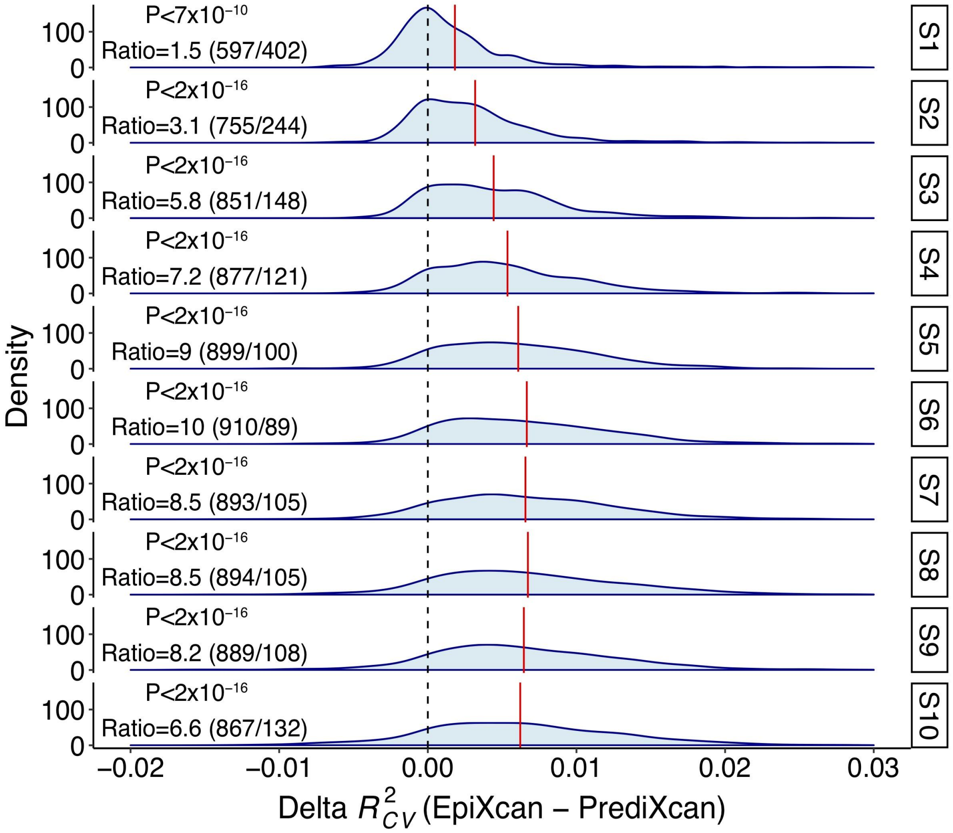

**Supplementary Figure 2. Simulation results.** Using simulated genotypes and gene expression in 500 samples, we compare the adjusted cross-validation (CV) $R^2$ of EpiXcan and PrediXcan by estimating the delta (Δ) $R^2_{CV}$ value (EpiXcan $R^2_{CV}$ minus PrediXcan $R^2_{CV}$). We simulated 10 scenarios, where in each scenario we increase the level of noise in the gene expression data. Across all simulations, the overall delta value is positive indicating that EpiXcan outperforms PrediXcan. *P* value from one-sample sign test is provided to compare whether the shift of the $\Delta R^2_{CV}$ values is different than zero ($H_0$: $\tilde{X} = 0$). The numbers in parentheses indicate the occasions where $\Delta R^2_{CV}$ was higher in EpiXcan ($\Delta R^2_{CV} > 0$; left number) and PrediXcan ($\Delta R^2_{CV} < 0$; right number); ratio is estimated by dividing the occasions of $\Delta R^2_{CV} > 0$ with $\Delta R^2_{CV} < 0$. The red vertical line shows the mean of delta value.

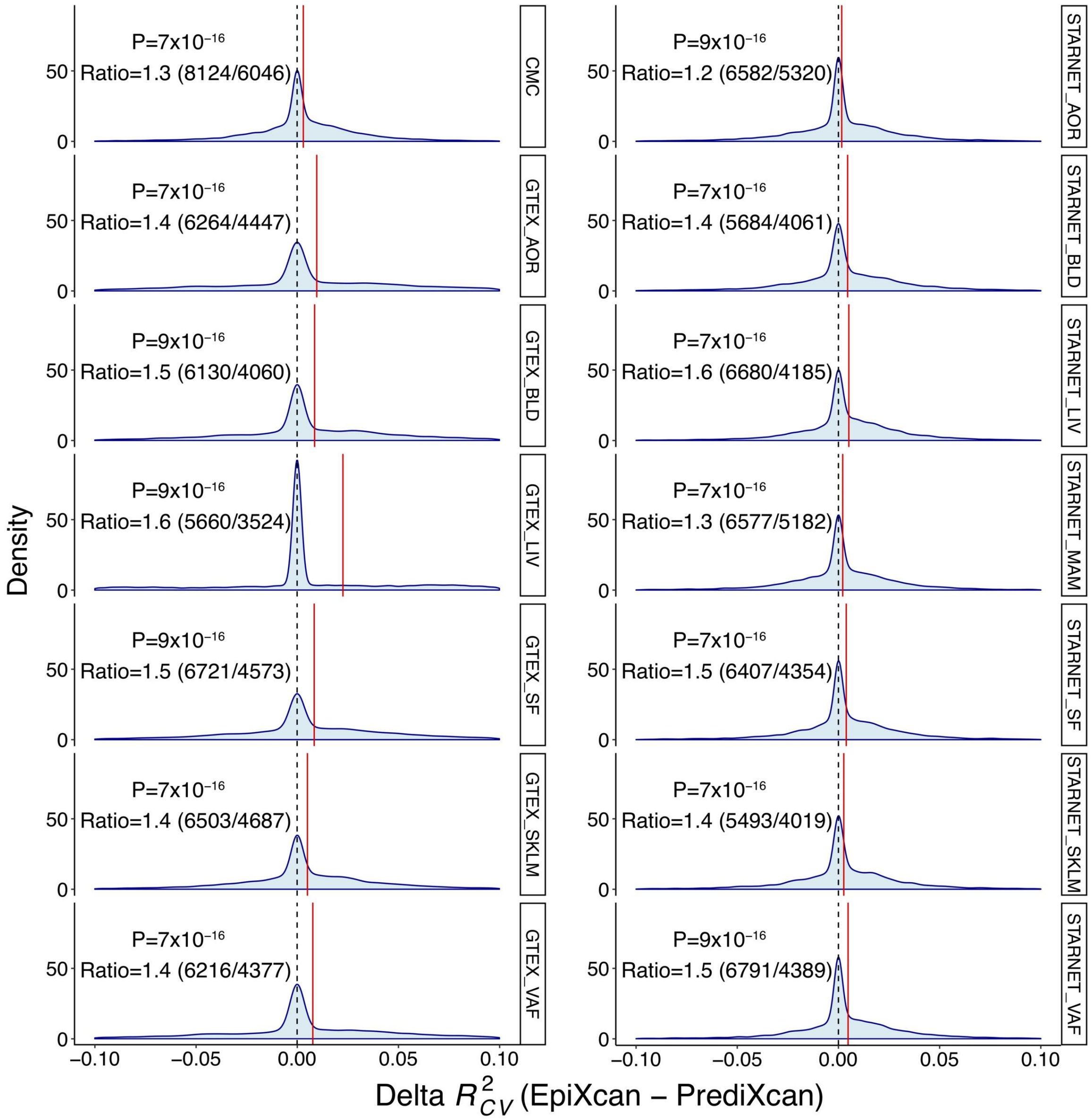

**Supplementary Figure 3. Comparison of model performance during training (cross validation).** We apply EpiXcan and PrediXcan in 14 tissue datasets and compare the adjusted $R^2_{CV}$ by estimating the $\Delta R^2_{CV}$ (EpiXcan $R^2_{CV}$ minus PrediXcan $R^2_{CV}$). Across all datasets, the overall delta value is positive indicating that EpiXcan outperforms PrediXcan. *P* value from one-sample sign test is provided to compare whether the shift of the $\Delta R^2_{CV}$ values is different than zero ($H_0$: $\tilde{X} = 0$). The numbers in parenthesis indicate the occasions where $\Delta R^2_{CV}$ was higher in EpiXcan ($\Delta R^2_{CV} > 0$; left number) and PrediXcan (delta $R^2_{CV} < 0$; right number); ratio is estimated by dividing the occasions of $\Delta R^2_{CV} > 0$ with $\Delta R^2_{CV} < 0$. The red vertical line shows the mean of delta value.

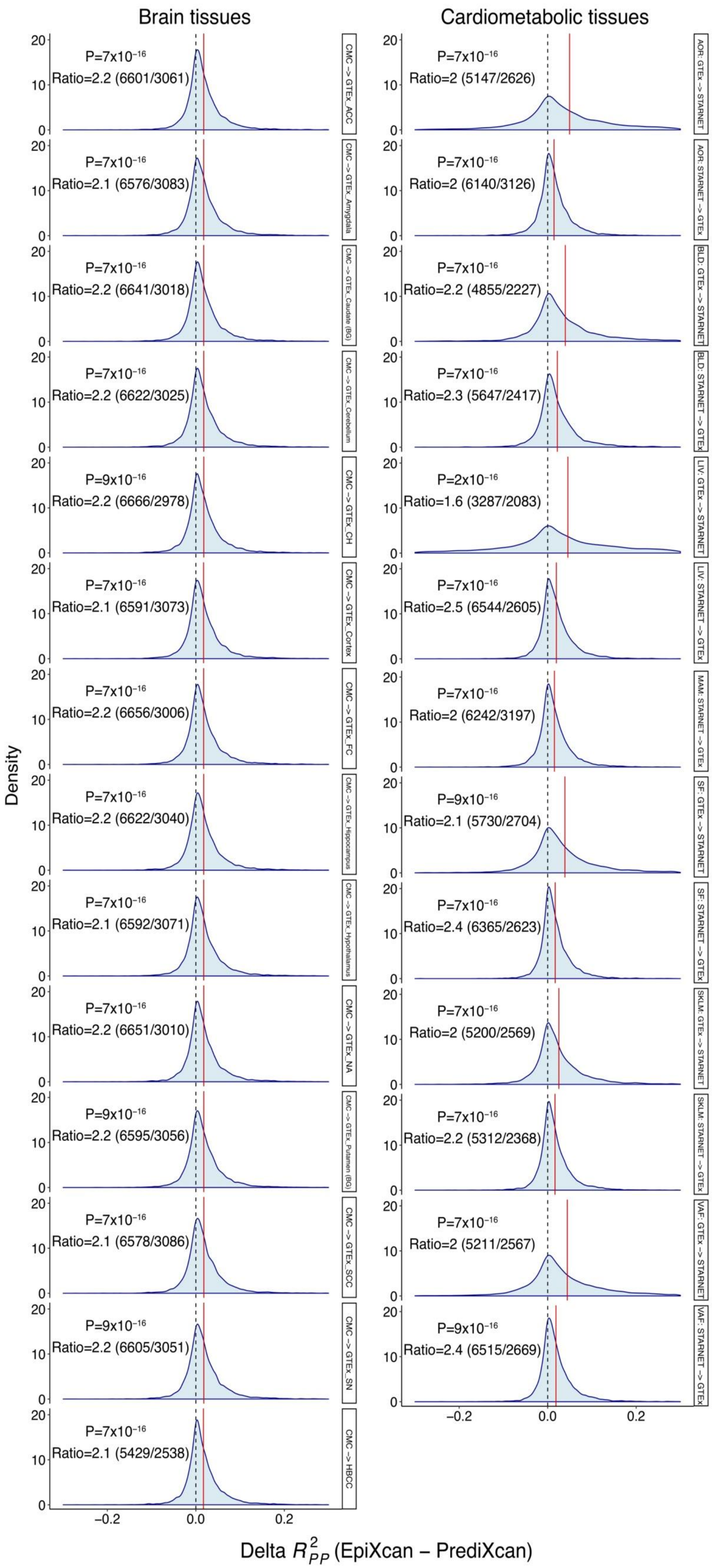

**Supplementary Figure 4. Comparison of model performance during prediction in independent datasets.** We used EpiXcan and PrediXcan models to predict expression levels in relevant brain and cardiometabolic independent datasets. We compare the adjusted predictive performance ($R^2_{PP}$) by estimating the $\Delta R^2_{PP}$ (EpiXcan $R^2_{PP}$ minus PrediXcan $R^2_{PP}$). Across all datasets, the overall delta value is positive indicating that EpiXcan outperforms PrediXcan. P value from one-sample sign test is provided to compare whether the shift of the $\Delta R^2_{PP}$ values is different than zero (H₀: $\tilde{X} = 0$). The numbers in parenthesis indicate the occasions where $\Delta R^2_{PP}$ is higher in EpiXcan ($\Delta R^2_{PP} > 0$; left number) and PrediXcan ($\Delta R^2_{PP} < 0$; right number); ratio is estimated by dividing the occasions of $\Delta R^2_{PP} > 0$ with $\Delta R^2_{PP} < 0$. The red vertical line shows the mean of delta value.

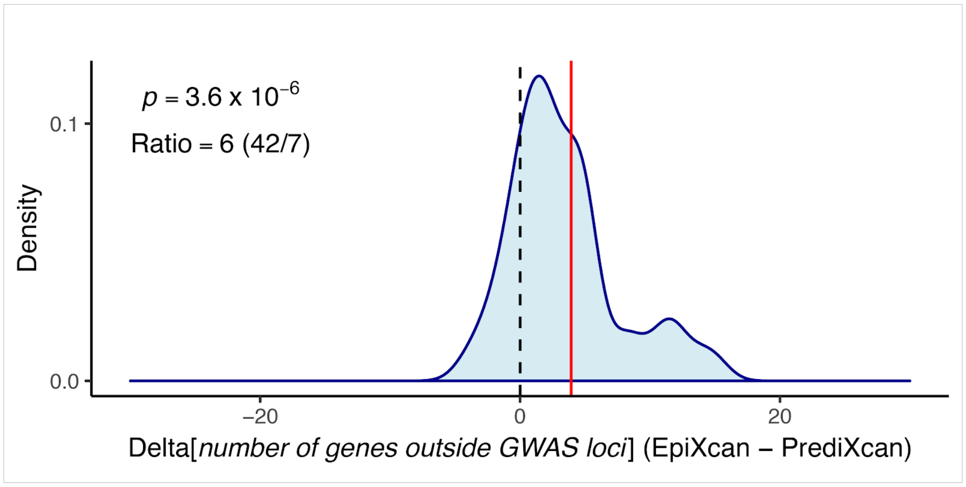

**Supplementary Figure 5. EpiXcan identifies more "novel genes" than PrediXcan.** The density plot, which shows the distribution of the Δ[number of genes outside GWAS loci] (EpiXcan – PrediXcan), shows that EpiXcan identifies more novel genes, which are within loci that did not reach genome-wide significance, than PrediXcan (one-sample sign test $p$ value = $3.6 \times 10^{-6}$, $H_0$: $\tilde{X} = 0$). Ratio is estimated by dividing the occasions of delta $n_{novel\ gene} > 0$ with delta $n_{novel\ gene} < 0$. The red vertical line shows the mean of delta values.

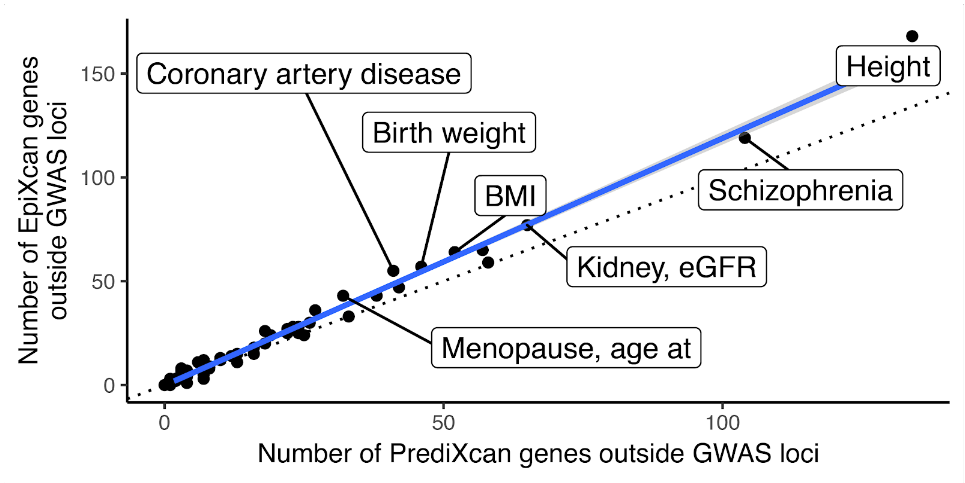

**Supplementary Figure 6. Number of "novel genes" identified by EpiXcan and PrediXcan.** From the scatter plot, we see that EpiXcan identifies more "novel genes" than PrediXcan per trait. The blue line corresponds to the regression line with 95% CI in grey. The dashed line is y=x.

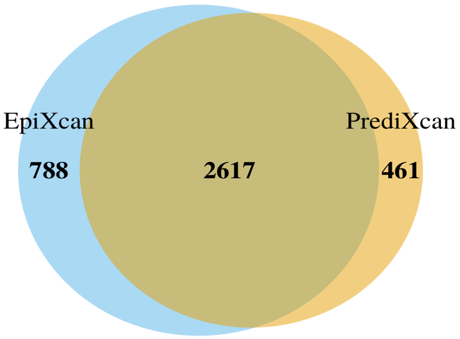

**Supplementary Figure 7. EpiXcan uniquely identifies more significantly associated genes than PrediXcan.** Venn diagram of genes with statistically significant gene-trait associations identified by both methods: EpiXcan and PrediXcan.

**a**

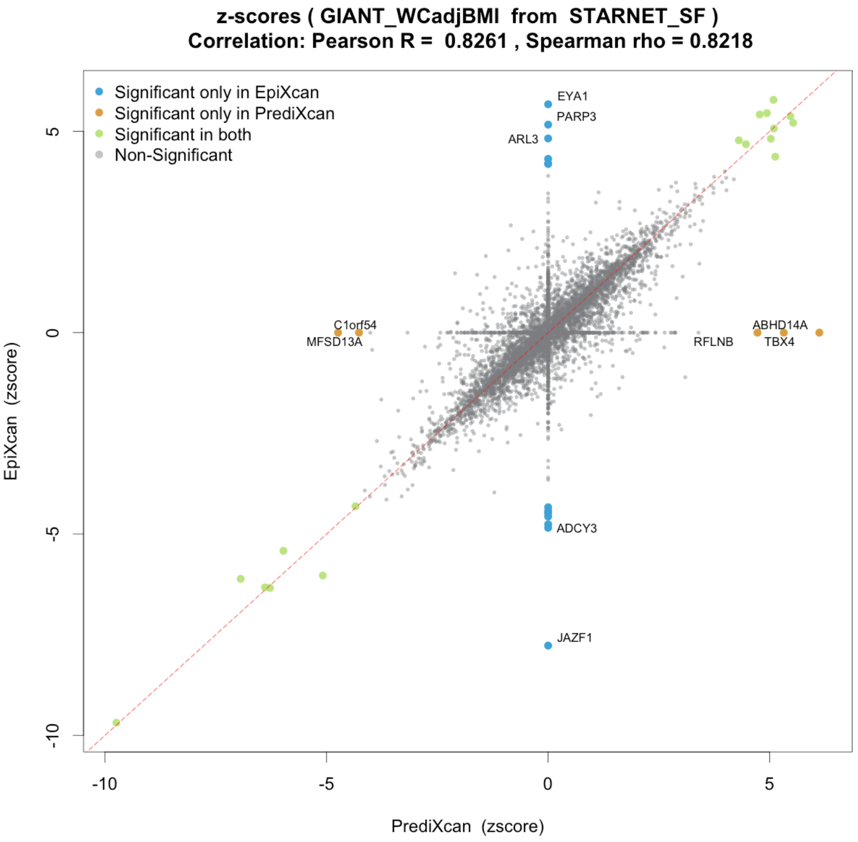

**b**

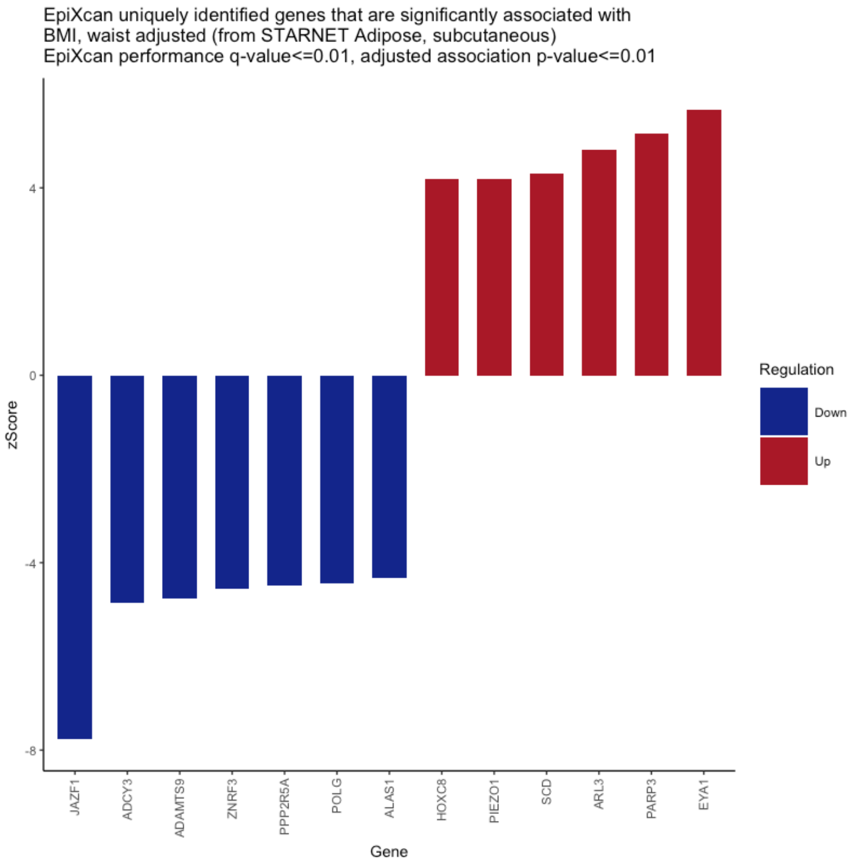

**Supplementary Figure 8. (a) Scatterplot demonstrating the correlation between z-score predictions by EpiXcan and PrediXcan for waist adjusted BMI in STARNET subcutaneous adipose tissue.** Grey dots indicate genes that are identified by both methods but are not significant (FDR>1%). All colored dots (blue, orange and light green) denote genes that significantly associated with the trait (waist circumference adjusted BMI). Green dots represent those significantly associated genes identified by both methods. Blue dots denote genes that are identified only by EpiXcan and orange dots those that are uniquely identified by PrediXcan; the top five genes based on [z] for each of the methods are named in the graph. Genes corresponding to uniquely identified genes by either method as above are not considered for the calculation of Pearson's and Spearman's correlation. **(b) Genes uniquely identified by EpiXcan for waist adjusted BMI in STARNET subcutaneous adipose tissue.** Z-scores of genes uniquely identified by EpiXcan corresponding to the blue dots in (a) panel. Respective (a) and (b) plots for all the 58 traits across the 14 tissues of the study can be found in our online repository http://icahn.mssm.edu/EpiXcan

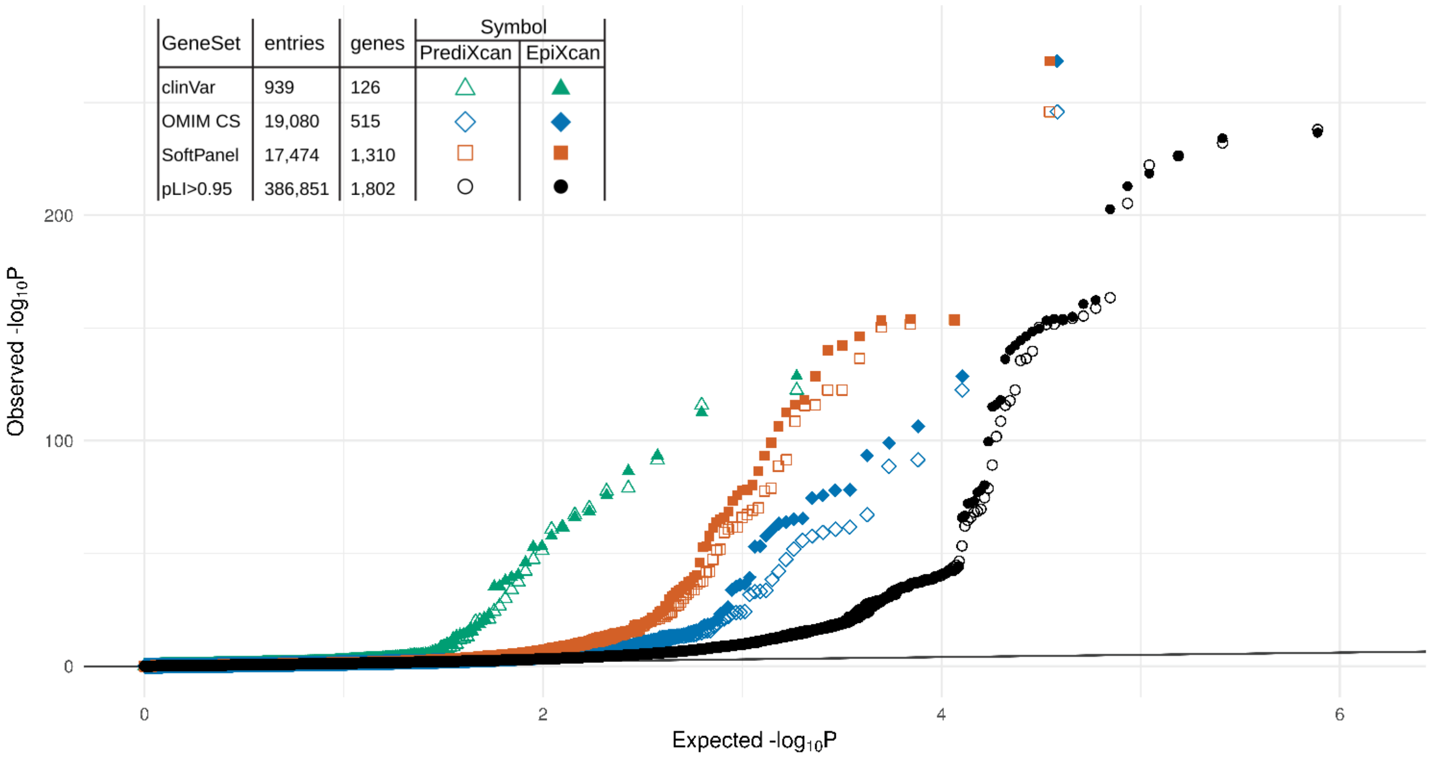

**Supplementary Figure 9. Enrichment of clinically significant genes in EpiXcan and PrediXcan gene-trait associations.** This is a Q-Q plot of the *p* values of the gene-trait associations for both EpiXcan and PrediXcan. For each human phenotype dataset (clinVar - green triangles, OMIM CS – blue diamonds, SoftPanel – orange squares) the entries are plotted for each gene, for all tissues but only for the traits for which the gene is associated in the respective dataset. In contrast, for the pLI > 0.95 dataset all points are plotted for all traits and tissues since there is not trait-specific information resulting to a higher number of entries. Since each entry represents a unique combination of gene, tissue (n=14) and trait (n=58), one gene can have up to 812 entries. All Q-Q plots are statistically significantly different (Kolmogorov-Smirnov against all values – not shown).

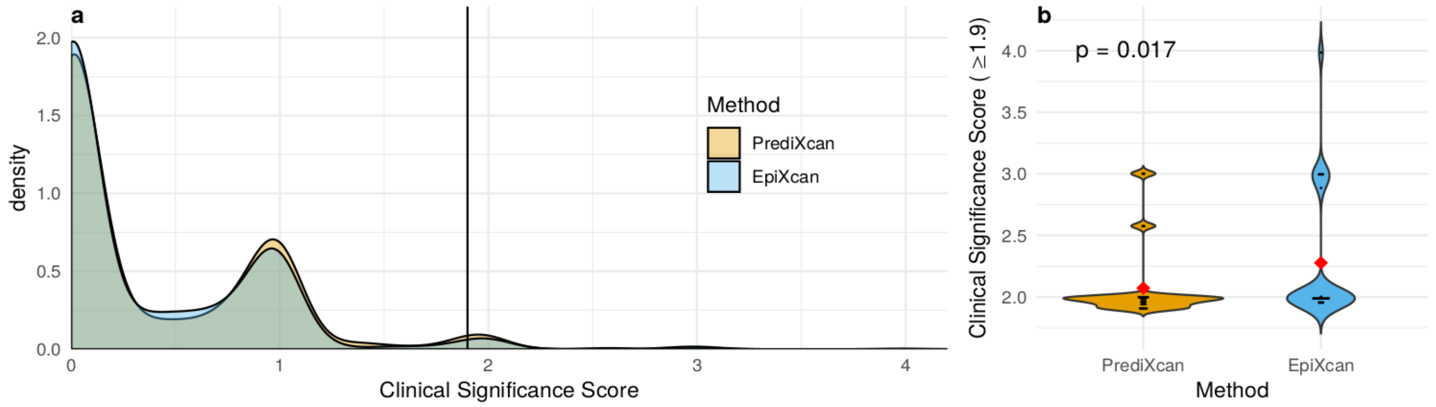

**Supplementary Figure 10. Uniquely identified genes by EpiXcan are more likely to have clinical significance as verified by more than one clinical significance datasets.** We define a clinical significance score (CSS) that ranges from 0 to 5, presence of a known gene-trait association in either of the datasets (ClinVar, OMIM CS, SoftPanel, MGD) counts for 1 point and then the pLI (0 to 1) is added to form the final score. (a) Density plot for the CSS of unique genes identified from EpiXcan and PrediXcan. The vertical line corresponds to CSS of 1.9 - the minimum score for a gene to have a gene-trait specific association corroborated by more than one datasets (eg. OMIM CS and pLI ≥ 0.9). (b) Violin plot for gene-trait associations with CSS ≥ 1.9. The red diamonds correspond to the mean of each distribution. P value was estimated with two-group Mann-Whitney U test.

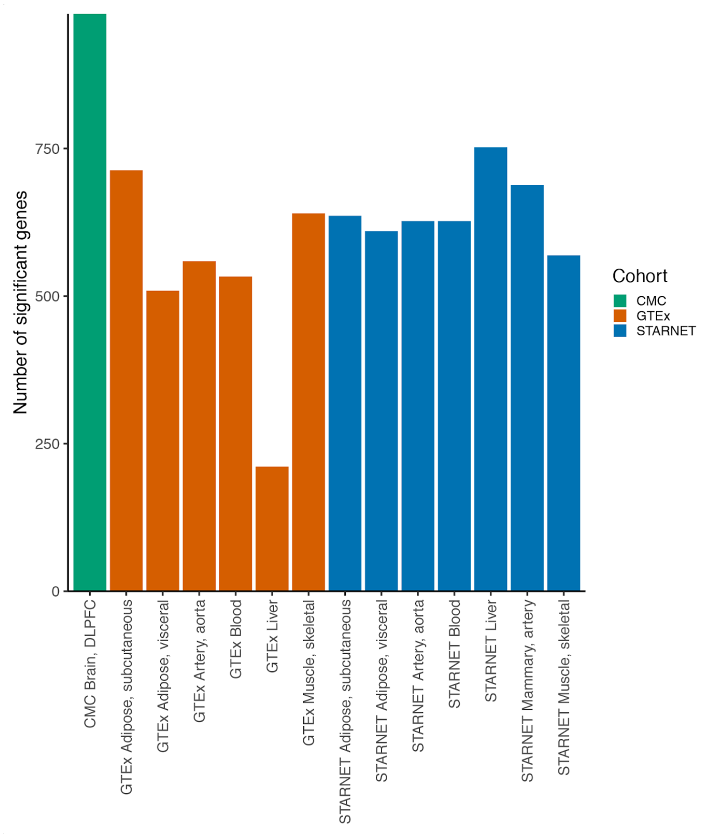

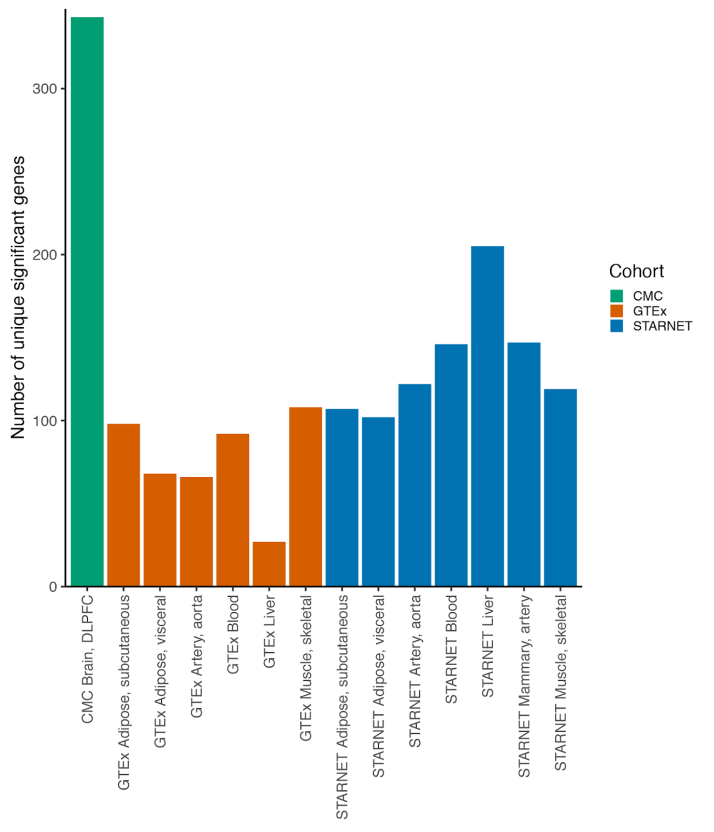

**a** **b**

**Supplementary Figure 11. Gene contributions from different tissue models**. (a) Contribution of significantly associated genes from each tissue in different cohorts. In total, there are 3,405 significant genes that identified for all traits. (b) Contribution of unique (as in only identified for the trait in this specific tissue) significant genes from each tissue/cohort. GTEx liver tissue contributes less than others, which is reasonable, due to the smallest sample size (n=130). For the size of studies, please refer to **Supplementary Table 1**.

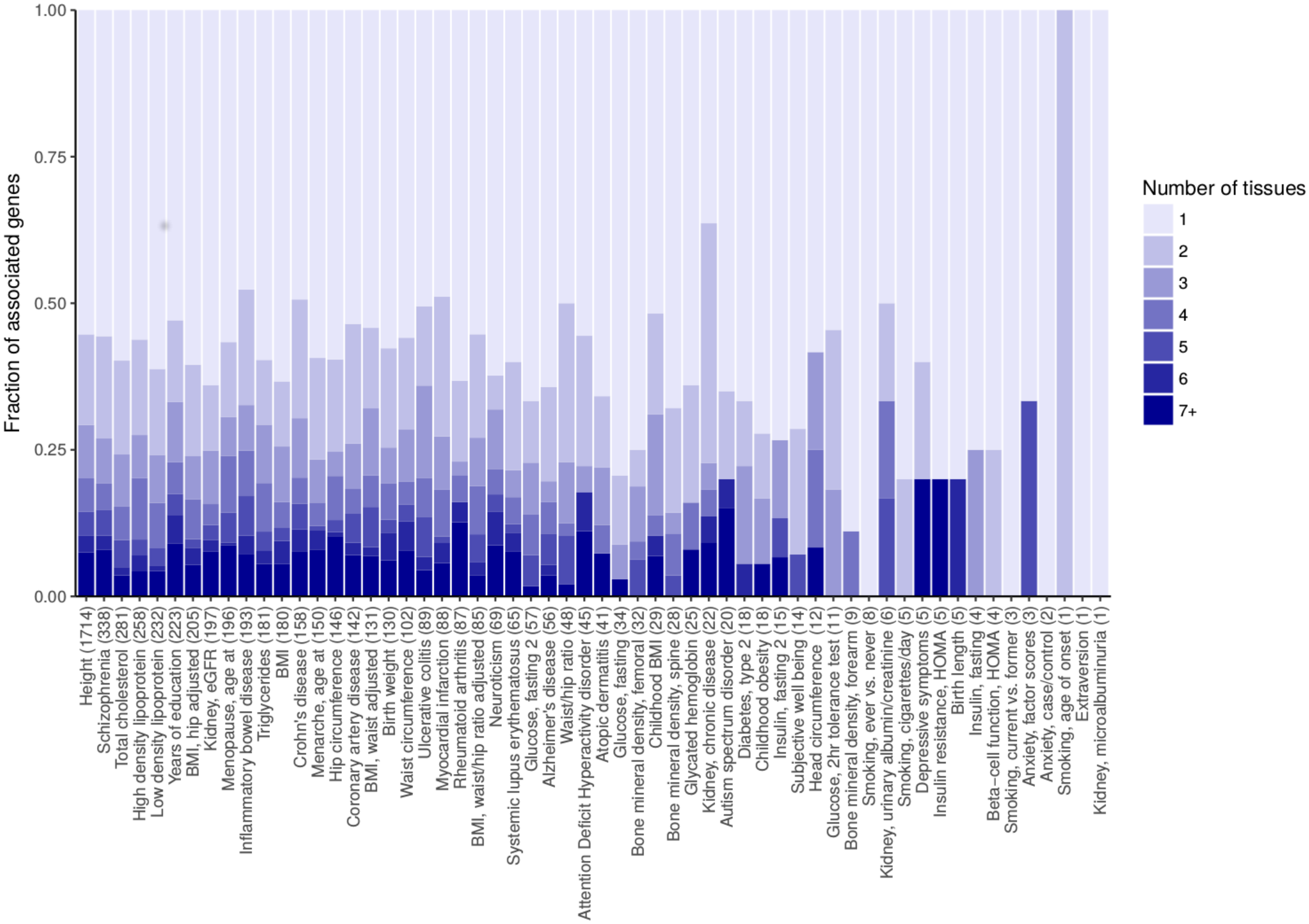

**Supplementary Figure 12. Proportions of number of tissues contributing trait-specific gene trait associations.** For each trait we show the fraction of associated genes that are contributed by a single tissue up to 7 or more tissues. Numbers in parentheses provide the total number of genes associated with each trait. We see that for most of the traits, more than 50% of the gene-trait associations are contributed from a single tissue.

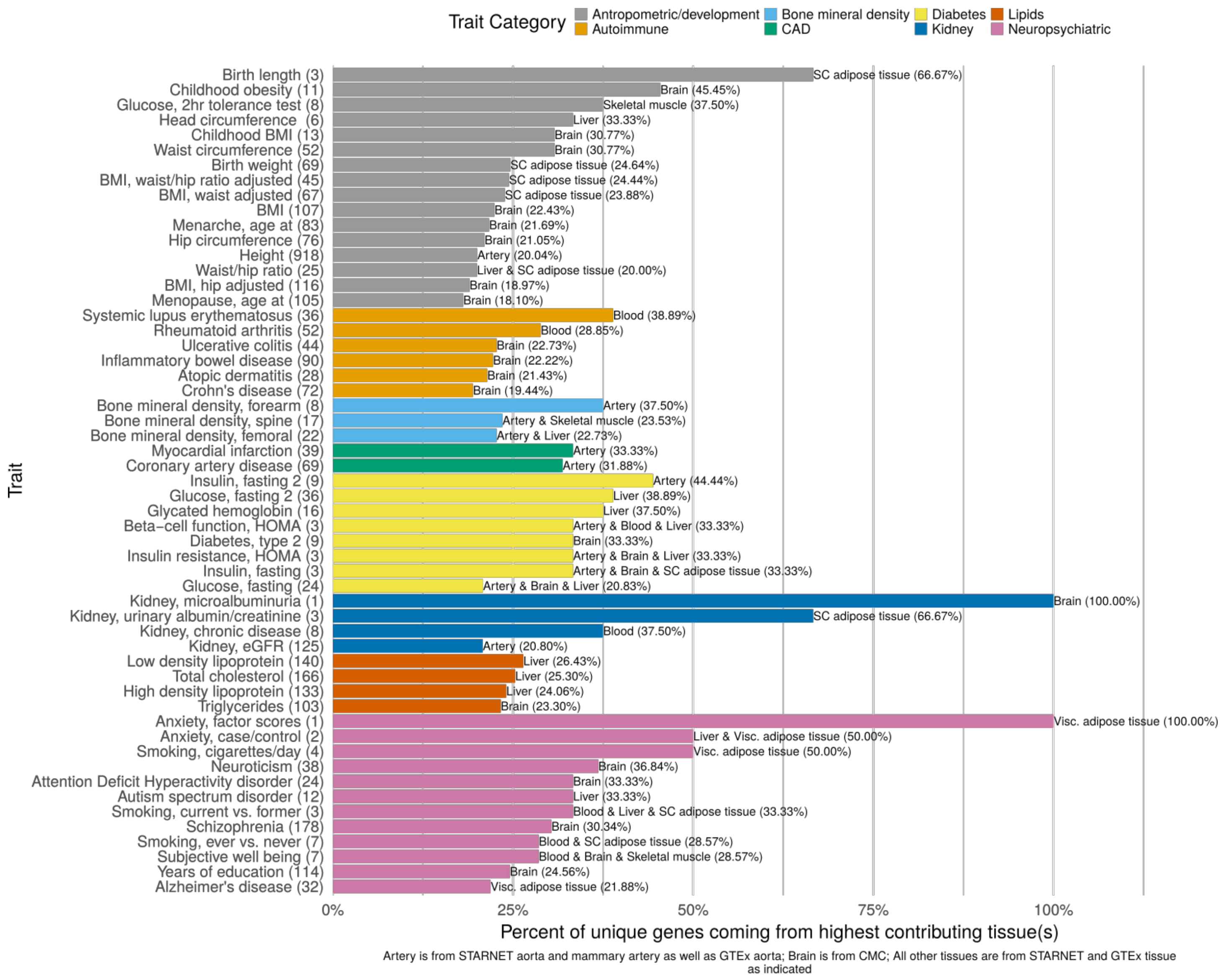

**Supplementary Figure 13**. **Percent of gene-trait associations contributed by top tissue type for each trait.** For almost all the traits, a very big proportion of the unique associated genes come from one tissue type (32.98% ± 17.36%; mean ± SD). The digits in the parentheses indicate the number of genes being contributed by only one tissue type. The bars denote percentages of unique genes coming from highest contributing tissues for all the traits. If more than one tissue contributes the same top number of unique genes, all tissue type names are provided (separated by "&").

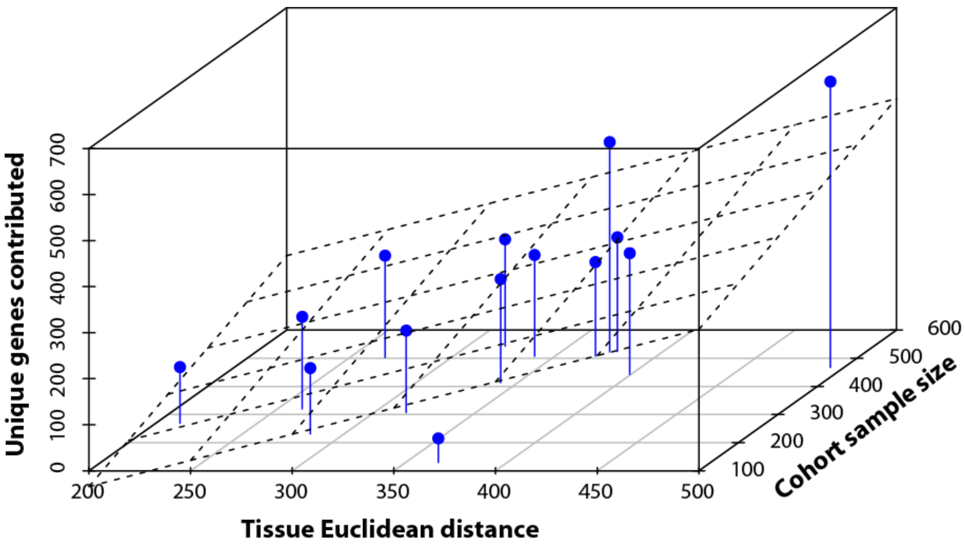

**Supplementary Figure 14. Cohort sample size and tissue dissimilarity explain most of the variation in the number of unique genes contributed by each tissue model.** 3D scatter plot demonstrating how tissue dissimilarity (as estimated by the average Euclidean distance of the significant $z$ scores versus all other tissues) and cohort sample size correlate with the number of unique genes contributed. Each blue dot represents one of the 14 tissue models of the study, the blue line projections (for each tissue i: $\langle x, y, z\rangle = \langle x_i, y_i, z_i\rangle + t\langle 0, 0, -z_i\rangle$) help create sense of depth for visualization. The plane corresponding to the multiple linear regression model ($N_{\text{unique genes}}$ = -300.78 + 1.13 × "Tissue Euclidean distance" + 0.39 × "Cohort sample size", adjusted $R^2 = 0.52$, $p$ value = 0.007) is drawn with dotted lines.

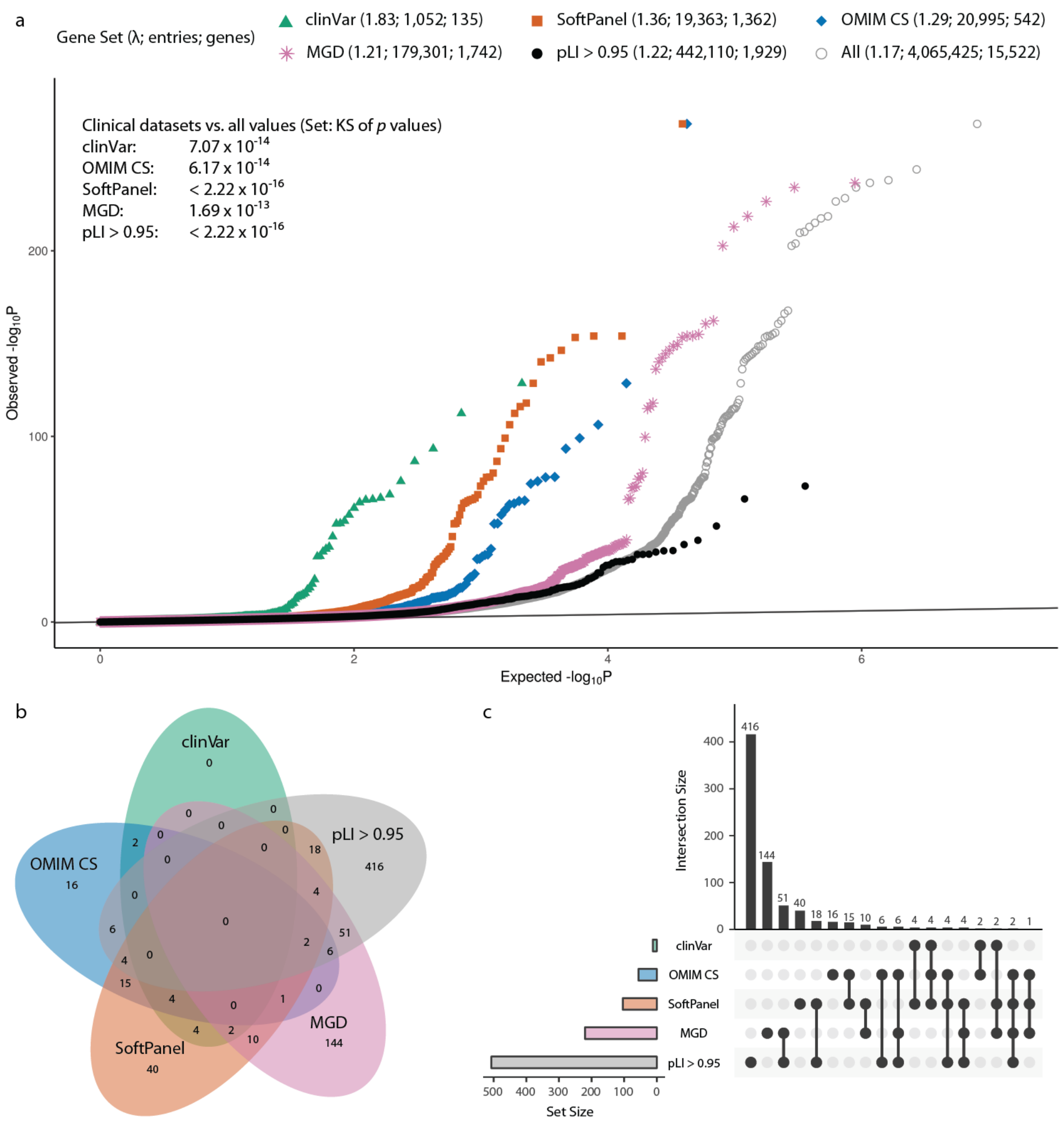

**Supplementary Figure 15. Enrichment of associated clinically relevant genes in EpiXcan gene-trait associations. (a) Q-Q plot of the *p* values of the gene-trait associations for EpiXcan.** For human phenotype datasets (**Supplementary Information**; clinVar - green triangles, OMIM CS – blue diamonds, SoftPanel – orange squares) the entries are plotted for each gene, for all tissues but only for the traits for which the gene is associated in the respective dataset; phenotypic severity is lower in OMIM CS which identifies clinical signs similar to the trait and higher in clinVar which corresponds most of the times to a Mendelian (monogenic) form of the trait. The entries are plotted similarly for the MGD dataset (MGD – pink stars) which identifies ortholog mouse genes that are associated with mouse phenotypes that are in the same broad phenotypic category as the human trait (**Supplementary Information**). In contrast, for the pLI > 0.95 dataset (black circles) all points are plotted for all traits and tissues since there is not trait-specific information. For reference, the *p* value distribution of all predictions is given (grey circles). Since each entry represents a unique combination of gene, tissue (n=14) and trait (n=58), one gene can have up to 812 entries. Genomic inflation factors (λ) are given in the legend at the top and Kolmogorov-Smirnov *p* values (against distribution of all values) are given in the custom annotation (top left). **Venn diagram (b) and matrix layout (c) for all the intersections of genes with statistically significant gene-trait associations that belong to at least one of the datasets in (a).**

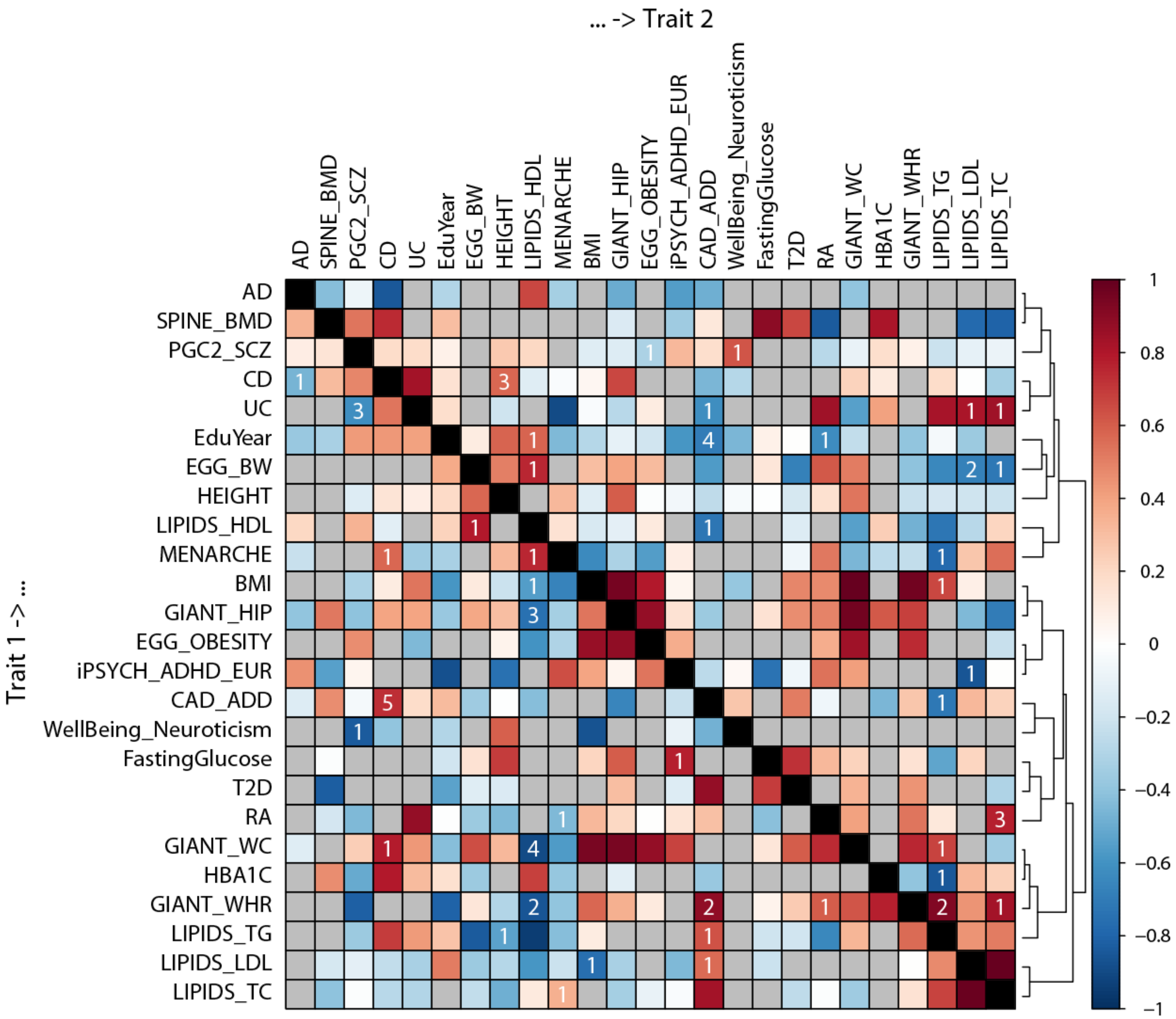

**Supplementary Figure 16**. **Putative trait causal relationships based on imputed transcriptomes.** Bi-directional regression analysis was performed for the predicted transcriptomes of all tissues for all significantly correlated trait pairs ($r_g$ and $r_{GReX}$, $q$ value ≤ 0.05) and the conditional estimates $\rho_{\text{Trait 1 | Trait 2}}$ are shown as color-coded squares in this 2D matrix (blue = protective = -1; red = causal = 1). Trait pairs that are not significantly correlated are shown with grey. For the trait pairs that displayed a significant difference (p<0.05, Welch's t test) in their conditional estimates (Trait 1 -> Trait 2 *vs.* Trait 2 -> Trait 1), we use white labels to denote the number of tissues where this difference was observed. Dendrogram on the right edge is shown from Ward hierarchical clustering.

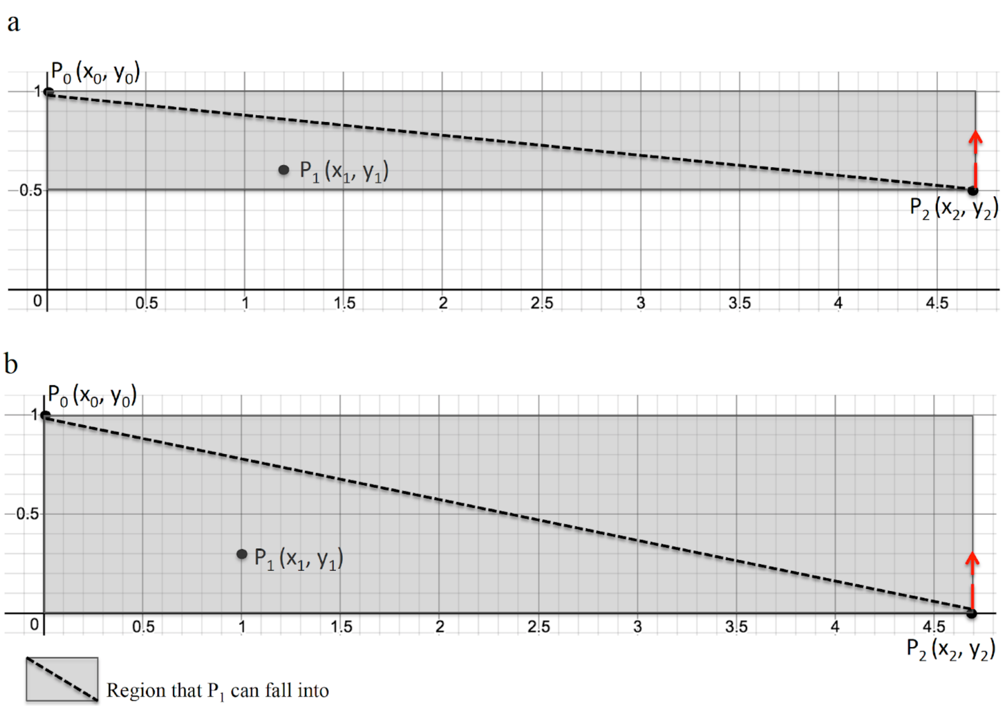

**Supplementary Figure 17. Demonstration of the process to determine the rescaling function by second order Bézier approximation.** By using 2nd order Bézier curve, we need to decide only one optimal intermediate control point $P_1$. $P_0$ is (0, 1), and $P_2$ has coordinate ($x_2$, $y_2$), where $x_2$ is the maximal value of all the priors and $y_2$ is fixed in each window. Red dashed arrow points to the direction of shifting the window. (a) The window of the process where $y_2 = 0.5$. (b) The first window of the process where $y_2=0$.

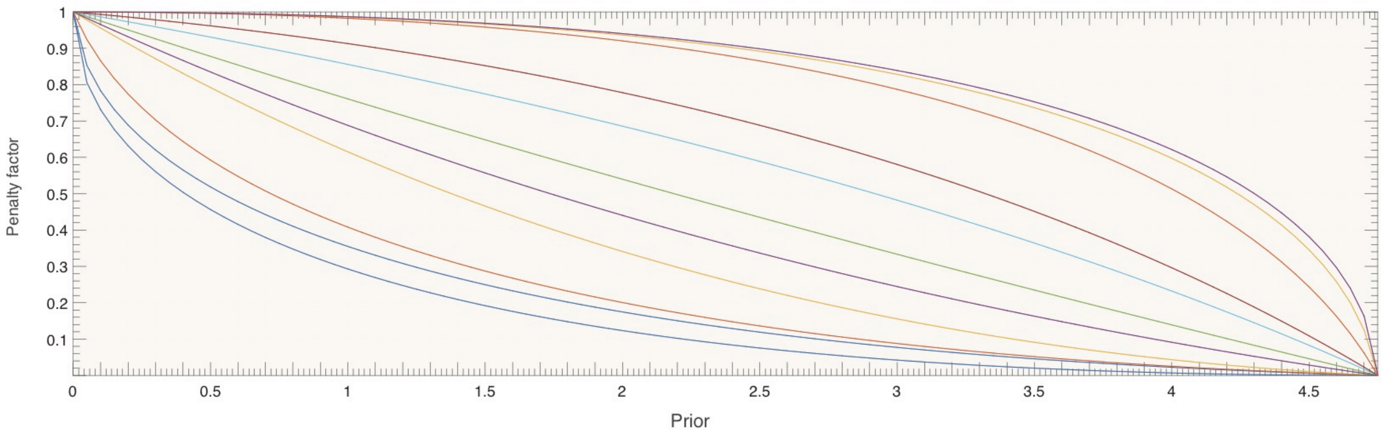

**Supplementary Figure 18. Examples of candidate rescaling's in one illustrative window.** The starting and end control points determine the start and end of the curve, but the intermediate control point determines the shape of the rescaling function. Every grid point in the window is one candidate intermediate control point, based on which different candidate interpolations are obtained. Since we use quadratic (second order) interpolation functions the direction of the curve can only change once thus creating either concave down or concave up curves in this set of examples. Higher order Bézier interpolation functions may have combination of concave down and concave up shapes.

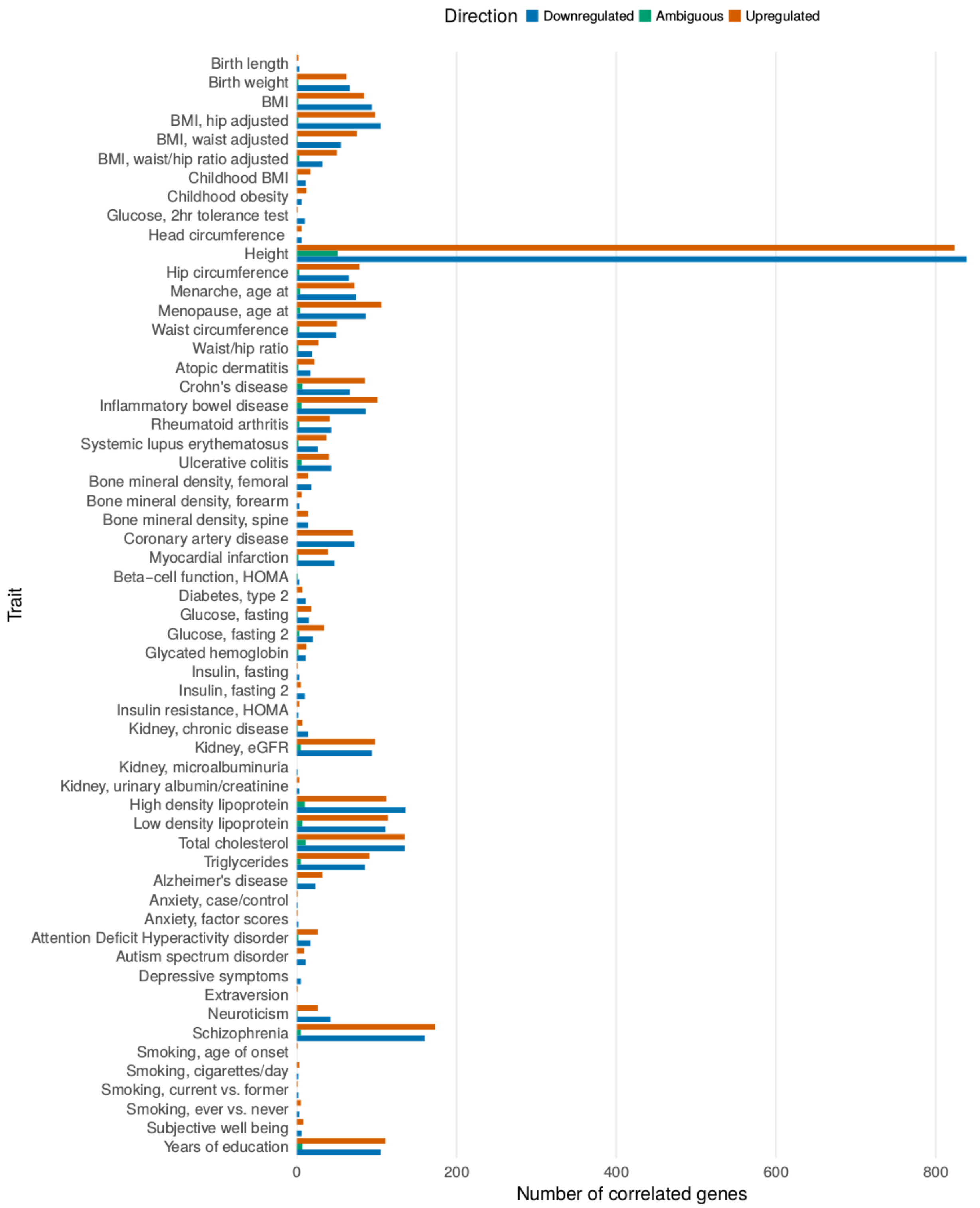

**Supplementary Figure 19. Genes predicted to be up- or down-regulated in different tissues for each of the traits.** A gene is regarded as up-/down-regulated for a given trait if z-score are positive/negative in the majority of the tissues. If there are equal numbers of tissues predicting up- or down-regulation, then the expressional change is regarded as ambiguous. Only genes that are significantly associated with traits are considered.

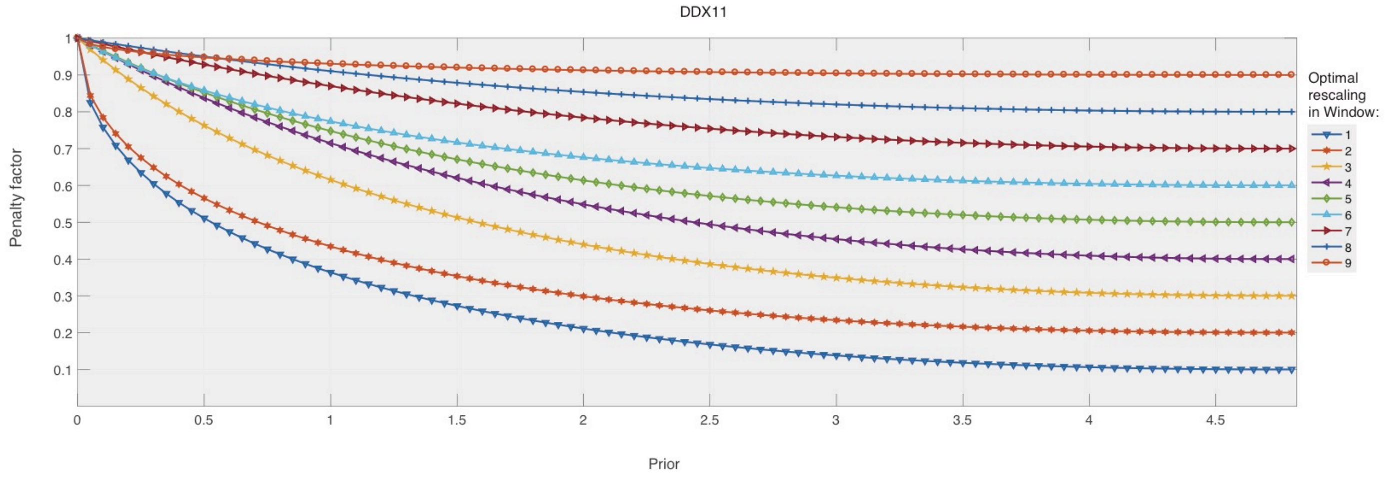

**Supplementary Figure 20. Optimal rescalings in each of the windows for *DDX11* simulations.** The rescaling in window 7 was found to be the best performing one.

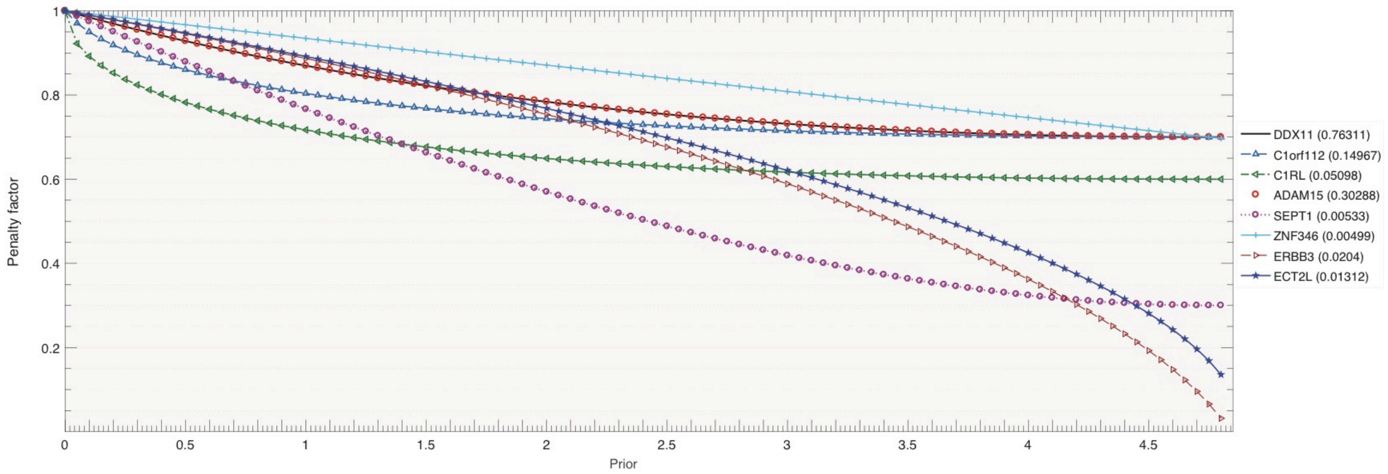

**Supplementary Figure 21**. **Optimal rescaling's regarding different simulations.** This plot shows the best rescaling for each of the simulations regarding several genes. Numbers in brackets indicate the $R^2_{CV}$ of the genes. Then the rescaling's are evaluated by the overall $R^2_{CV}$ and the mapping(s) from *DDX11* (*ADAM15*) is assessed to be the best optimal rescaling. The mappings from these two genes are exactly the same and more important, the overall $R^2_{CV}$ of the rescaling is superior to others. Overall $R^2_{CV}$ is calculated based upon the real CMC data. For each rescaling, we compare the overall $R^2_{CV}$ with that of PrediXcan using Wilcoxon test, and the rescaling with most significant *p* value is considered as most optimal.

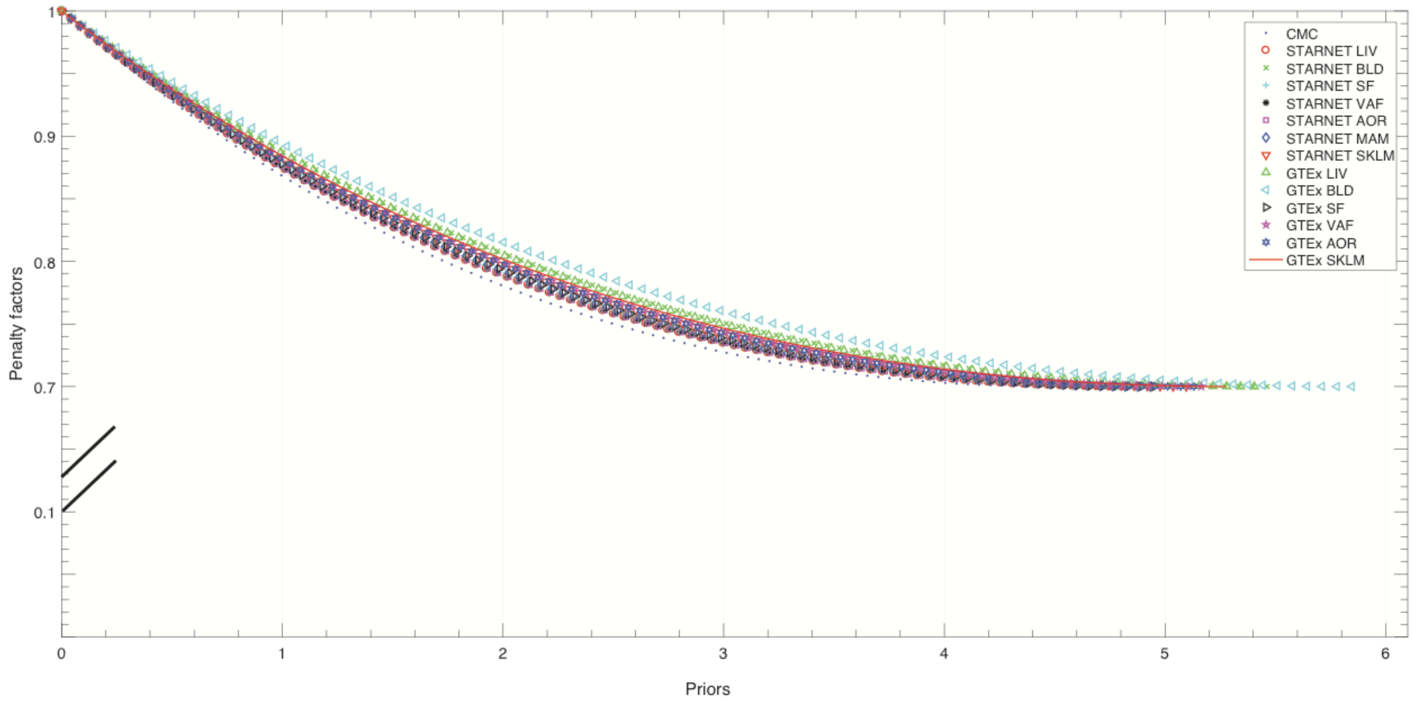

**Supplementary Figure 22**. **Optimal rescaling's for each of the data sets.** Due to different prior ranges of each data set, the rescaling functions vary.

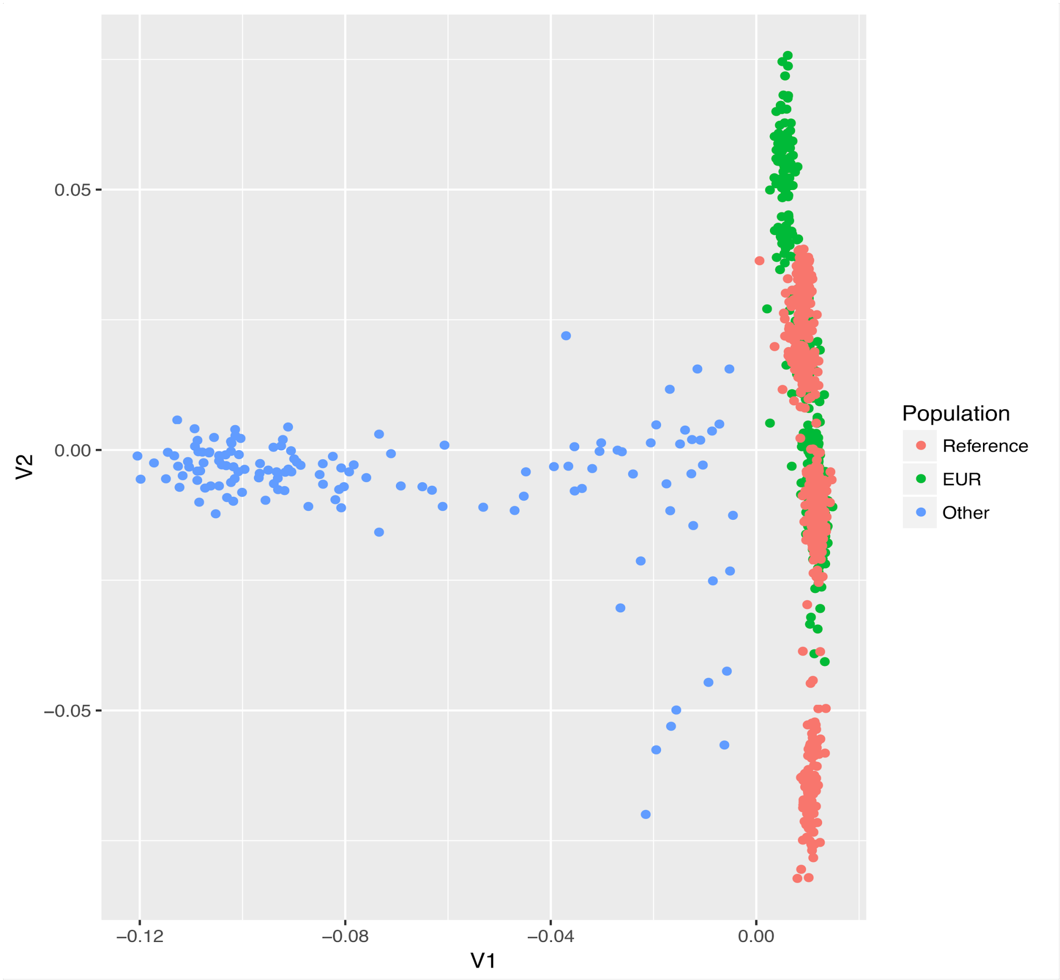

**Supplementary Figure 23. Principle analysis to identify European population for CMC.**

### Legends for Supplementary tables

**Supplementary Table 1. Overview of gene expression and genotype datasets included in current study.** "Number of Genes" is the number of genes that are detectable in each dataset. "Predictor" indicates the datasets used to train PrediXcan and EpiXcan models. "Observed" indicates the datasets used to verify accuracy of predictions.

**Supplementary Table 2. Comparison of $R^2_{CV}$ between EpiXcan approach and PrediXcan.** *p* values show significances of EpiXcan $R^2_{CV}$ improvements over PrediXcan $R^2_{CV}$ using Wilcoxon pair-wise test with 'greater' option.

**Supplementary Table 3. Comparison of prediction correlations, $R^2_{PP}$, between EpiXcan and PrediXcan.**

**Supplementary Table 4. Information of the 58 traits and GWAS datasets.** In total, there are 58 GWASs that considered. This table gives the resource and information of each GWAS that used in our analysis. Corresponding full names of the traits are listed and it also provides the category that every trait belongs to. We give the number of genes that significantly associate with each trait. The numbers of genes that up-/down-regulated with all of the traits are listed.

**Supplementary Table 5. GSEA - pLI (excel file with 4 sheets).** Gene set enrichment analysis (GSEA) for all pLI (probability of loss of function intolerant) deciles. We first test for enrichment among all significant genes identified by each method (sheets: "EpiXcan_all" and "PrediXcan_all" respectively). Then we test for enrichment among genes specific to each trait categories identified by each method (sheets: "EpiXcan_trait_categories" and "PrediXcan_trait_categories" respectively"). GSEA is performed for all pLI deciles, p values are calculated with the fisher exact test and are FDR-adjusted to q values.

**Supplementary Table 6. Pathway analysis of significantly associated genes in 43 traits.** In order to study if the associated genes are specific to biological processes regarding various diseases, we performed gene-set enrichment analysis of 43 traits with >10 significantly associated genes. After corrections, 74 highly enriched pathways are obtained with $P < 1.70\times10^{-5}$, and adjusted $P < 0.0488$. This table provides top pathways in which the associated genes are enriched.

**Supplementary Table 7. Computational drug repurposing hits for eQTL traits, and chemogenomic enrichment analysis.**

**Supplementary Table 8. Trait association pairs.** For each pair of traits, we calculate the number of significantly associated genes that they share. Only traits that associated with at least 50 genes are considered. The number of significantly associated genes of each trait is listed. In total, there are 311 association pairs, 245 of which are crossing trait categories. 66 association pairs are within trait categories.

**Supplementary Table 9. qtlBHM priors of each dataset.** This table lists the prior values of SNPs within every annotation region that integrated.

**Supplementary Table 10. Query parameters used to infer gene-trait associations from the clinical datasets.**
